# A gated hydrophobic funnel within BAX binds long-chain alkenals to potentiate pro-apoptotic function

**DOI:** 10.1101/2024.12.23.630122

**Authors:** Jesse D. Gelles, Yiyang Chen, Mark P. A. Luna-Vargas, Ariele Viacava Follis, Stella G. Bayiokos, Jarvier N. Mohammed, Tara M. Sebastian, M. Abdullah Al Noman, Ngoc Dung Pham, Yi Shi, Richard W. Kriwacki, Jerry E. Chipuk

**Affiliations:** Laboratory of Mitochondrial Biology in Human Health and Disease, Icahn School of Medicine at Mount Sinai, One Gustave L. Levy Place, New York, New York 10029, USA; Department of Oncological Sciences, Icahn School of Medicine at Mount Sinai, One Gustave L. Levy Place, New York, New York 10029, USA; The Tisch Cancer Institute, Icahn School of Medicine at Mount Sinai, One Gustave L. Levy Place, New York, New York 10029, USA; The Graduate School of Biomedical Sciences, Icahn School of Medicine at Mount Sinai, One Gustave L. Levy Place, New York, New York 10029, USA; Department of Structural Biology, St. Jude Children’s Research Hospital, 262 Danny Thomas Place, Memphis, Tennessee 38105, USA; Department of Pharmacological Sciences, Icahn School of Medicine at Mount Sinai, One Gustave L. Levy Place, New York, New York 10029, USA; Department of Microbiology, Immunology and Biochemistry, University of Tennessee Health Sciences Center, Memphis, Tennessee 38105, USA; Department of Dermatology, Icahn School of Medicine at Mount Sinai, One Gustave L. Levy Place, New York, New York 10029, USA; The Diabetes, Obesity, and Metabolism Institute, Icahn School of Medicine at Mount Sinai, One Gustave L. Levy Place, New York, New York 10029, USA

**Keywords:** α, β-Unsaturated Alkenals, Apoptosis, BAX, BCL-2 Family, Cell Death, Hexadecenal, MOMP

## Abstract

Mitochondria maintain a biochemical environment that cooperates with BH3–only proteins (e.g., BIM) to potentiate BAX activation, the key event to initiate physiological and pharmacological forms of apoptosis. The sphingosine-1-phosphate metabolite 2-trans-hexadecenal (2t–hexadecenal) is one such component described to support BAX activation, but molecular mechanisms remain largely unknown. Here, we utilize complementary biochemical and biophysical techniques to reveal that 2t-hexadecenal non-covalently interacts with BAX, and cooperates with BIM to stimulate early-activation steps of monomeric BAX. Integrated structural and computational approaches reveal 2t–hexadecenal binds an undefined region – a hydrophobic cavity formed by core-facing residues of α5, α6, and gated by α8 – we now term the “BAX actuating funnel” (BAF). We define alkenal length and α8 mobility as critical determinants for 2t–hexadecenal synergy with BIM and BAX, and demonstrate that proline 168 allosterically regulates BAF function. Collectively, this work imparts detailed molecular insights advancing our fundamental knowledge of BAX regulation and identifies a regulatory region with implications for biological and therapeutic opportunities.

## INTRODUCTION

Developmental, homeostatic, and pharmacological pro-apoptotic signals converge by engaging the BCL-2 family of proteins to induce BAX-dependent mitochondrial outer membrane permeabilization (MOMP) and apoptosis.^1^ Despite sequence and structural similarity with multiple globular BCL-2 family proteins, BAX is unique in that it converts from an inactive cytosolic monomer to pore-forming oligomer at the outer mitochondrial membrane (OMM). Upon transient triggering with direct activator BH3–only proteins (e.g., BIM, BID), BAX undergoes a series of intramolecular rearrangements and structural refoldings that ultimately result in translocation to the OMM, oligomerization, and MOMP.^2–7^ An important structural event during this process is α9 helix mobilization from its residence within the BAX BC groove, which simultaneously supports OMM translocation, BAX:BAX interactions, and propagation of the activation process leading to proteolipid pore formation. We refer to this series of structural rearrangements as the BAX activation continuum, which may be separated into an activation phase (triggering) and functionalization phase (pore formation).^8^

There are two general requirements for potent BAX activation: protein-protein interactions to trigger the monomer, and protein-lipid interactions to initiate and stabilize refolding of BAX into multimeric conformers. These requirements are proposed to occur in distinct phases of the activation continuum, and most insights into protein-lipid interactions focus on how BAX interacts within the OMM to form pore-like structures.^9–12^ This has led to a modern conceptualization in which mitochondrial membranes actively cooperate with the BCL–2 family to control cell death commitment.^13^ Notably, mitochondrial endoplasmic reticulum contact sites (MERCS) maintain lipid homeostasis between the organelles and supply lipids that are critical for MOMP.^14^ One example, the terminal end-product of the sphingolipid pathway – 2-trans-hexadecenal (2t–hexadecenal) – is required for BAX-mediated MOMP following triggering by BIM or BID.^15^ 2t–hexadecenal is formed by the irreversible cleavage of sphingosine-1-phosphate (S1P) and is metabolized to palmitoyl-CoA for fatty acid synthesis pathways.^16^ The resident enzymes responsible for generating and catabolizing 2t–hexadecenal are enriched at MERCS, and genetically modulating 2t–hexadecenal levels alters sensitivity to human BAX in yeast.^17^ Additionally, interactions between BAX and the OMM are governed by OMM curvature, which is regulated by a combination of the mitochondrial dynamics machinery, intra-organellar membrane contact sites, and mitochondria-specific lipids.^18,19^

While the requirement for 2t–hexadecenal in the BAX activation continuum is described, cooperation with BH3–only proteins and underlying mechanisms remain poorly understood. Here, we utilize biochemical, biophysical, structural, and computational approaches to systematically demonstrate that 2t–hexadecenal directly actuates BAX through non-covalent interactions at a previously undefined region – a funnel-shaped hydrophobic cavity formed by core-facing residues of α5, α6, and gated by α8, which we term the BAX Actuating Funnel, or BAF. Our results suggest that BIM-mediated BAX triggering mobilizes α8, making the BAF accessible, and that binding of 2t–hexadecenal promotes BAX functionalization. Furthermore, we identify chemical and structural determinants underlying the 2t–hexadecenal:BAX interaction and reveal that mutation of proline 168 in the loop between α8 and α9 allosterically deforms the BAF and subsequently disrupts the function of 2t–hexadecenal. Collectively, this model advances our understanding of the BAX structure-function relationship by characterizing the protein-lipid interactions responsible for stimulating monomeric BAX activation and identifies a previously underappreciated regulatory domain for both cell biology and therapeutic investigations.

## RESULTS

### 2t–hexadecenal directly activates BAX through non-covalent interactions

Previous work demonstrated that hexadecenal was required for potent BAX-mediated MOMP and that sphingolipid precursors are supplied to mitochondria via interactions with heterotypic membranes.^15^ To assess whether ectopic hexadecenal exposure could engage BAX activation *in cellulo*, we treated SV40-transformed mouse embryonic fibroblasts (MEFs) with increasing concentrations of hexadecenal and measured the apoptotic response using our real-time multi-visitation microscopy technique, SPARKL.^20^ Supraphysiological concentrations of ectopic hexadecenal (“2t–16”) did not induce cell death at the lower concentrations tested (Figure 1A). We reasoned that any pro-apoptotic signaling may have been mitigated by the repertoire of anti-apoptotic BCL–2 family proteins, and indeed, co-treatment with the BH3-mimetic ABT-737 revealed an apoptotic phenotype in response to ectopic hexadecenal (Figure 1A). To determine if the apoptotic response required triggering by direct activators, we utilized *Bim^−/−^Bid^−/−^* double knockout (DKO) MEFs and observed a loss of apoptosis at lower concentrations of ectopic hexadecenal co-treated with ABT-737, suggesting that BAX activation was part of the underlying mechanism (Figure 1B). Finally, we replicated this experiment in *Bax^−/−^Bak^−/−^* DKO MEFs and observed no cell death (Figure 1C). Collectively, these data support the conclusion that hexadecenal induces apoptotic cell death by acting on effector BCL–2 family proteins.

**Figure 1:**
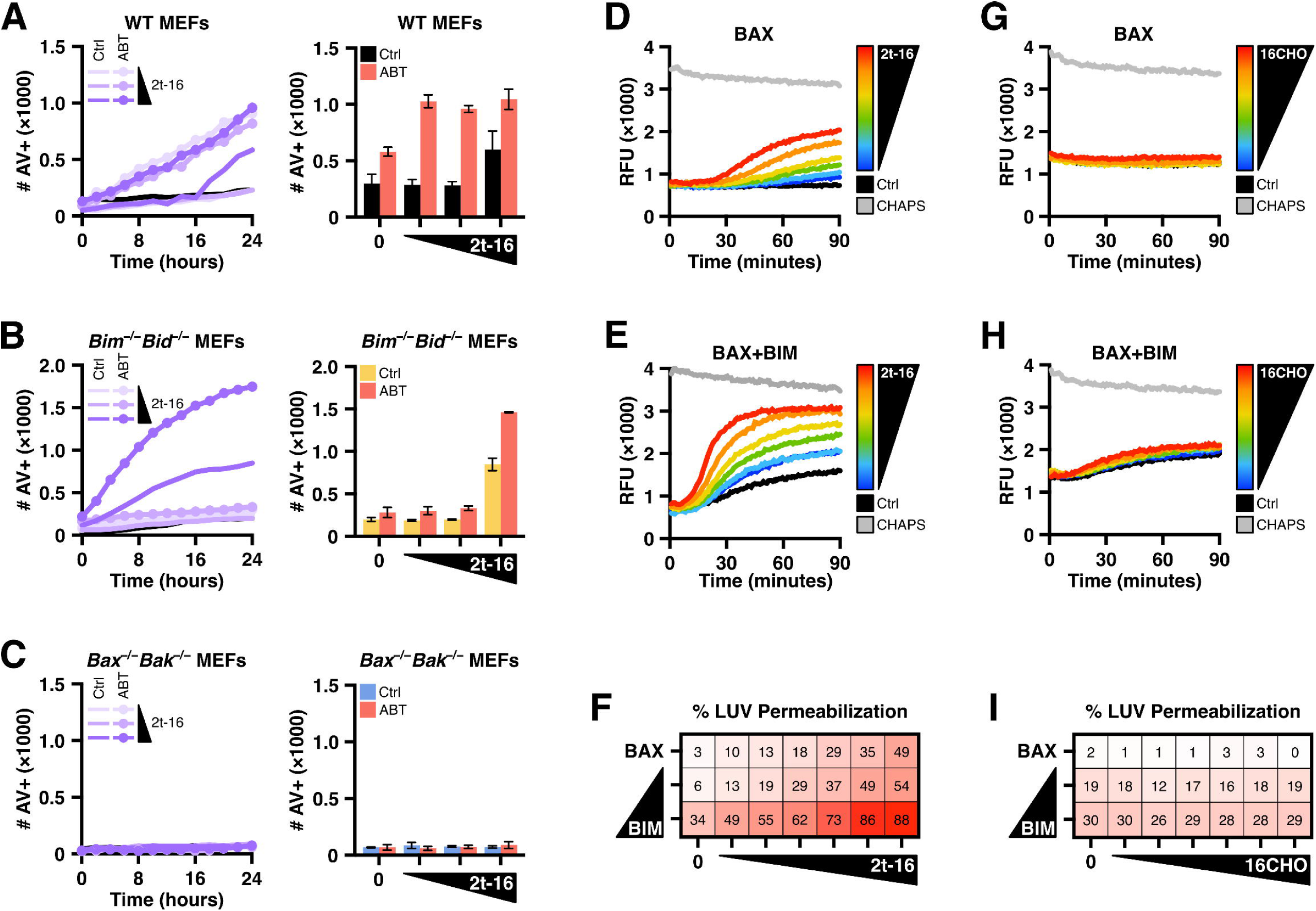
2t-hexadecenal induces apoptosis and membrane permeabilization by cooperating with BAX. **(A–C)** MEFs subjected to SPARKL analysis measuring real-time labeling with fluorescently-tagged Annexin V (100 μg/ml) via imaging with an IncuCyte ZOOM. Left panels: kinetics of cell labeling in response to increasing concentrations of 2t–hexadecenal (2t–16) with DMSO vehicle or co-treated with ABT-737 (ABT); black line reports untreated control. Right panels: endpoint data of replicates at 24 hours. Data shown are the mean of technical triplicates and error bars report SEM. **(A)** WT MEFs (matched to *Bax^−/−^Bak^−/−^* double knockout MEFs) were treated with 2t–16 (10, 20, 40 μM) and DMSO or ABT–737 (1 μM), imaged every 2 hours, and quantified for number of Annexin V-positive objects. **(B)** Same as in **A** with *Bim^−/−^Bid^−/−^* double knockout MEFs. **(C)** Same as in **A** with *Bax^−/−^Bak^−/−^* double knockout MEFs. **(D–I)** LUV permeabilization studies with recombinant BAX protein treated as indicated and measured at regular intervals for changes in fluorescence as fluorophores are released from liposomes. Grey data report LUVs solubilized with 1% CHAPS to measure maximal signal. Data shown are the mean of technical replicates. **(D)** BAX protein (120 nM) was combined with DMSO vehicle or 2t–16 (6.5–50 μM) followed by addition of LUVs and measured by fluorescent spectroscopy. **(E)** Same as in **D** with BIM–BH3 peptide (2.5 μM) added to BAX and 2t–16. **(F)** Heatmap visualization of normalized endpoint LUV permeabilization data from LUVs incubated with BAX (120 nM) treated with 2t–16 (6.5–50 μM) ± BIM–BH3 peptide (0.5, 2.5 μM). Data summarized from Figures **S1C**. **(G–H)** LUV permeabilization studies as in **D–E** with BAX (160 nM) and hexadecanal (6.5–50 μM) ± BIM–BH3 peptide (2.5 μM). **(I)** Heatmap visualization of normalized endpoint LUV permeabilization data as in **F** with BAX (160 nM) and 16CHO (6.5−50 μM). Data summarized from Figures **S1D**. See also Figure S1.

There are reports that ectopic hexadecenal can form adducts with DNA and generate oxidative stress resulting in apoptosis^21,22^; therefore, we interrogated whether hexadecenal directly promoted BAX-mediated pore formation by utilizing recombinant BAX protein and large unilamellar vesicles (LUVs), which are biochemically-defined liposomes that mimic the major lipid composition of the OMM, and assessed BAX activation by measuring LUV permeabilization. While recent studies have incorporated hexadecenal directly into the formulation of LUVs^23^, we aimed to investigate hexadecenal-mediated BAX activation without altering the biochemistry of the LUV membrane. BAX treated with supraphysiological concentrations of hexadecenal demonstrated dose-dependent activation and LUV permeabilization (Figure 1D). Importantly, ectopic hexadecenal alone did not disrupt or cause leakage of LUVs (Figure S1B). In the cell, BAX activation is mediated primarily through BCL–2 family direct activators – predominantly, BIM^24^ – and we had observed inhibited apoptosis in the *Bim^−/−^Bid^−/−^* MEFs; therefore we sought to assess the cooperation of BIM and hexadecenal on BAX-mediated membrane permeabilization. We treated BAX with an activating concentration of BIM–BH3 peptide and observed a dose-dependent increase and acceleration in LUV permeabilization in response to hexadecenal (Figures 1E, S1A). Furthermore, the synergy with hexadecenal was observed with mildly-activating concentrations of BIM–BH3 as well (Figures 1F, S1C). Previous work indicated that the saturated form of 2t–hexadecenal – hexadecanal (“16CHO”) – did not induce BAX oligomers in cross-linking studies^15^, and indeed, we did not observe BAX-mediated pore formation or synergy with BIM in response to hexadecanal (Figures 1G–I, S1D). These data indicate that hexadecenal promotes BAX pore formation and synergizes with BIM activation.

Despite treating BAX with hexadecenal directly, we could not entirely rule out the possibility that increased permeabilization could have been due to the lipidic aldehyde interacting with LUVs and resulting in a more permissive environment for BAX pore formation. Therefore, we utilized microscale thermophoresis (MST) to determine whether hexadecenal directly bound to BAX and observed a dose-dependent shift indicating changes to the molecular volume of BAX in response to hexadecenal (Figure 2A). Interestingly, the saturated hexadecanal aldehyde did not induce a similar change in BAX thermophoresis, suggesting that the α,β double bond is necessary for BAX interaction and subsequent activation (Figures 2B, S2A).

**Figure 2:**
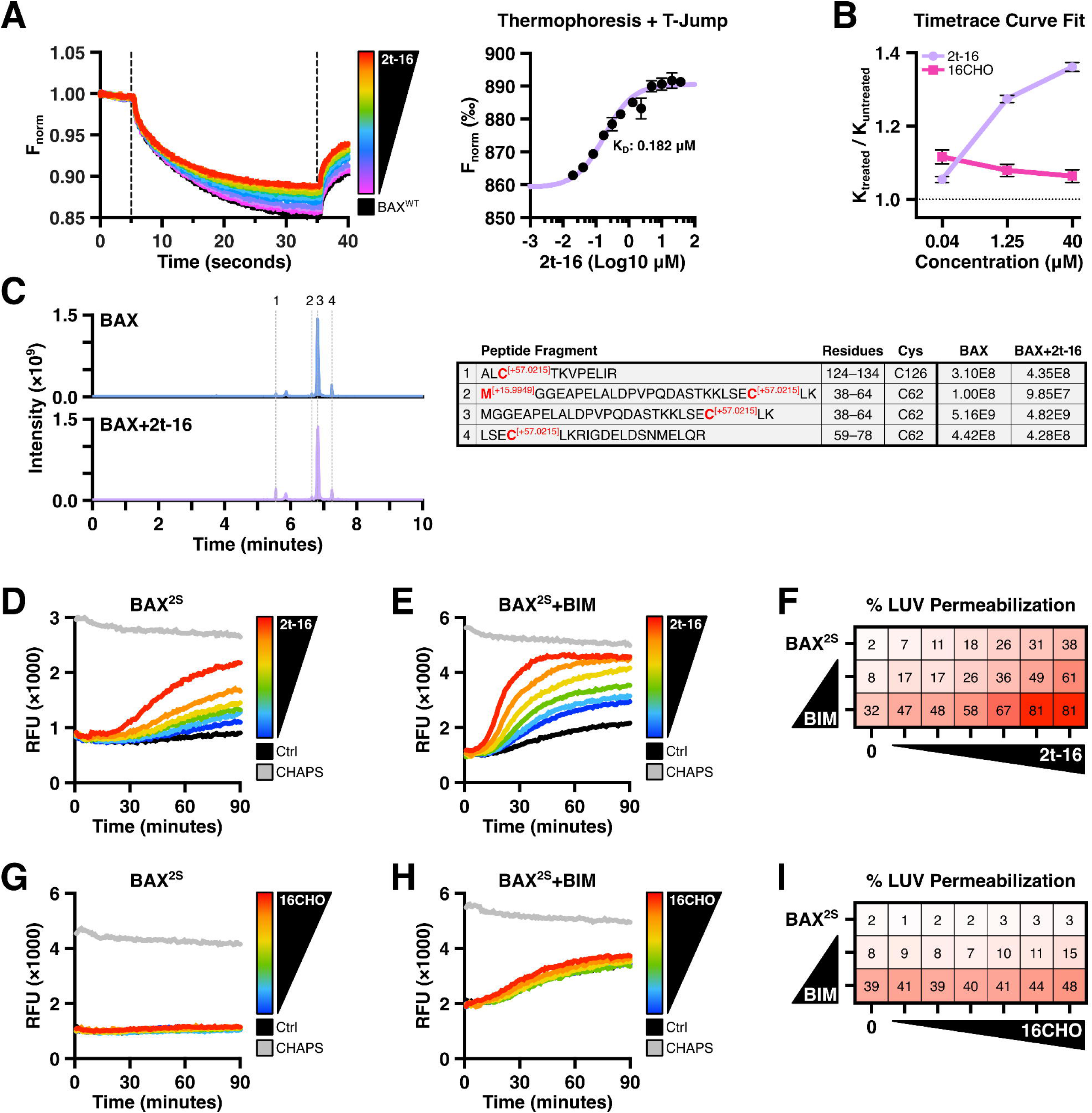
2t-hexadecenal activation of BAX is mediated by non-covalent interactions. **(A–B)** Alexa Fluor 647-labeled recombinant BAX^WT^ (1 nM) was incubated with CHAPS (0.002%) to inhibit oligomerization, treated as indicated, and subjected to MST. Data shown are the mean of replicate data and error bars report SD. **(A)** Left: Timetrace thermal shift curves of BAX^WT^ titrated with 2t–16 (0.02–40 μM) and subjected to MST. Right: Thermophoresis and temperature jump value for BAX^WT^ treated with a range of 2t–16 concentrations fitted to determine a K_D_ value. **(B)** BAX^WT^ was treated with 2t–16 or 16CHO (0.04, 1.25, 40 μM) and MST timetrace thermal shift curves were fitted using a one-step exponential function and compared using the decay (K) constants normalized by the untreated BAX curve. Original data in Figure **S2C**. **(C)** LC-MS of recombinant BAX^WT^ alone or incubated with 2t–16. Samples were then alkylated with iodoacetamide to identify unmodified cysteine residues and trypsin digested for analysis. Four cysteine-containing peptide fragments were detected. Values denote peptide abundance, calculated as AUC for each peak. **(D–I)** LUV permeabilization studies with recombinant BAX^2S^ protein treated as indicated and measured at regular intervals for changes in fluorescence as fluorophores are released from liposomes. Grey data report LUVs solubilized with 1% CHAPS to measure maximal signal. Data shown are the mean of technical replicates. **(D)** BAX^2S^ protein (100 nM) was combined with DMSO vehicle or 2t–16 (6.5–50 μM) followed by addition of LUVs and measured by fluorescent spectroscopy. **(E)** Same as in **D** with BIM–BH3 peptide (2.5 μM) added to BAX^2S^ and 2t–16. **(F)** Heatmap visualization of normalized endpoint LUV permeabilization data from LUVs incubated with BAX^2S^ (100 nM) treated with 2t–16 (6.5–50 μM) ± BIM–BH3 peptide (0.5, 2.5 μM). Data summarized from Figure **S2B**. **(G–H)** LUV permeabilization studies as in **D–E** with BAX^2S^ (100 nM) and hexadecanal (6.5–50 μM) ± BIM–BH3 peptide (2.5 μM). **(I)** Heatmap visualization of normalized endpoint LUV permeabilization data from LUVs incubated with BAX^2S^ (100 nM) treated with 16CHO (6.5–50 μM) ± BIM–BH3 peptide (0.5, 2.5 μM). Data summarized from Figure **S2C**. See also Figure S2.

Hexadecenal is an α,β-unsaturated aldehyde and is capable of modifying nucleophiles, (e.g., cysteine residues) through Michael addition, and the requirement for the double bond suggested a chemical reaction mechanism.^25^ In fact, recent publications have reported that hexadecenal covalently modifies BAX, though the studies disagree on which cysteine residue is modified and it is unclear whether the reaction is biologically specific to BAX.^23,26^ To determine whether the mechanism of hexadecenal-mediated BAX activation was the result of covalent modification, we incubated BAX with hexadecenal and subjected the sample to higher-energy collision-induced dissociation (HCD) and tandem liquid chromatography-mass spectrometry (LC-MS). We detected peptide fragments covering both cysteines (C62, C126) and observed no alkylation by hexadecenal (*m*+238.229 Da); as a control, we were able to detect modification of the cysteines by the alkylating agent iodoacetamide (*m*+57.021 Da) (Figure 2C). Additionally, we also observed no shift by intact mass spectrometry (data not shown). To decisively conclude that the mechanism was independent of cysteine modification, we replicated our LUV permeabilization studies using a cysteine-replacement BAX mutant (BAX^C62S,C126S^, “BAX^2S^”), which exhibited no changes to stability or melting temperature (Figure S2B). Compared to wild-type BAX (“BAX^WT^”), BAX^2S^ remained sensitive to hexadecenal-mediated changes in melting temperature, pore formation, and synergy with BIM–BH3 peptide (Figures 1D–F, 2D–F, S2C−D). Additionally, BAX^2S^ remained similarly unaffected by the saturated hexadecanal aldehyde (Figures 2G–I, S2E). Collectively, these results indicate that hexadecenal promotes BAX pore formation through a direct, non-covalent interaction mechanism.

### 2t–hexadecenal synergizes with BIM and promotes early-activation steps of monomeric BAX

The BAX activation continuum can be divided into two distinct phases of activation and functionalization: the cytosolic monomer gets activated and undergoes intramolecular rearrangements and conformational changes that result in translocation to the OMM; subsequently, active BAX proteins in the OMM undergo large-scale conformational changes, oligomerize, and mature into pore-forming units. Our data thus far measured BAX functionalization (i.e., pore formation) and demonstrated that hexadecenal cooperates with BIM to promote membrane permeabilization. Previous studies demonstrating hexadecenal-mediated BAX activation were mostly limited to model membranes with endogenous or incorporated hexadecenal, often at supraphysiological concentrations.^15,23^ In contrast, we utilized a direct treatment model of hexadecenal and BAX protein, indicating that the mechanism of action likely occurs on BAX found in the cytosol and prior to integration within the OMM. To investigate the effect of hexadecenal specifically on the activation phase of BAX, we utilized a technique we developed called FLAMBE, which monitors activation-induced intramolecular rearrangements within BAX (i.e., rearrangements that result in early-activation structural hallmarks such as displacement of the α1–α2 loop and mobilization of the C-terminal α9 helix).^4,8^ FLAMBE observes real-time early-activation of BAX by measuring changes in Polarization resulting from the kinetic binding and dissociation of a TAMRA-labeled BAK–BH3 peptide (BAK^TAMRA^); broadly, BAX activation can be inferred by measuring a reduction of Polarization over time (Figure S3A). Kinetic FLAMBE data can further be parameterized into time-of-maximum-Polarization (Tmax) and endpoint Polarization (EP) for trend overviews and comparative analyses between treatment conditions (Figures S3B). As an example, BAX treated with a range of BIM concentrations exhibits a dose-dependent pattern of BAK^TAMRA^ dissociation (measured as a reduction in Polarization over time) indicative of BAX activation (Figure S3C).

Using FLAMBE, we observed BAX^WT^ activation in the presence of hexadecenal as demonstrated by a reduction in Polarization over time, indicating that high concentrations of hexadecenal could induce a “direct-activator”-like effect on BAX monomers (Figure 3A). To confirm that this phenotype was not due to covalent modification of cysteine residues, we replicated the FLAMBE experiment with BAX^2S^ and observed the same activation profile (Figure 3B). By contrast, neither BAX^WT^ nor BAX^2S^ activated in response to the saturated hexadecanal aldehyde, though we did observe some destabilization of the BAX:BAK^TAMRA^ heterodimer at the higher concentrations (Figure S3D–E). We hypothesized that hexadecenal cooperates with direct activators instead of activating BAX *de novo* within the cell, and thus we investigated whether the synergy between hexadecenal and BIM occurs during the early-activation phase. We treated BAX^2S^ with a non-activating concentration of BIM–BH3 to generate a population of stable BIM:BAX:BAK^TAMRA^ heterotrimers (Figure 3C, yellow data). The addition of a non-activating concentration of hexadecenal resulted in activation of the primed BAX^2S^ population (Figure 3C, left panel); moreover, activation of primed BAX^2S^ was observed with several non-activating concentrations of hexadecenal (Figure 3C, right panel). When we induced a “triggered” BAX population (i.e., mildly activated by BIM), we similarly observed increased activation by non-activating concentrations of hexadecenal, as measured by increased kinetics of BAK^TAMRA^ dissociation, greater shifts in parameterized metrics, and reduction in area under the curve (Figures 3D, S3F). Importantly, we observed the same outcomes with BAX^WT^, confirming that the mechanism was not due to the endogenous cysteines or their mutation (Figure S3G). To eliminate the possibility that hexadecenal was inducing BAK^TAMRA^ dissociation by competing for the BC groove, we treated recombinant BCL–xL in the presence of BAK^TAMRA^ and observed no competition by hexadecenal (Figure S3H).

**Figure 3:**
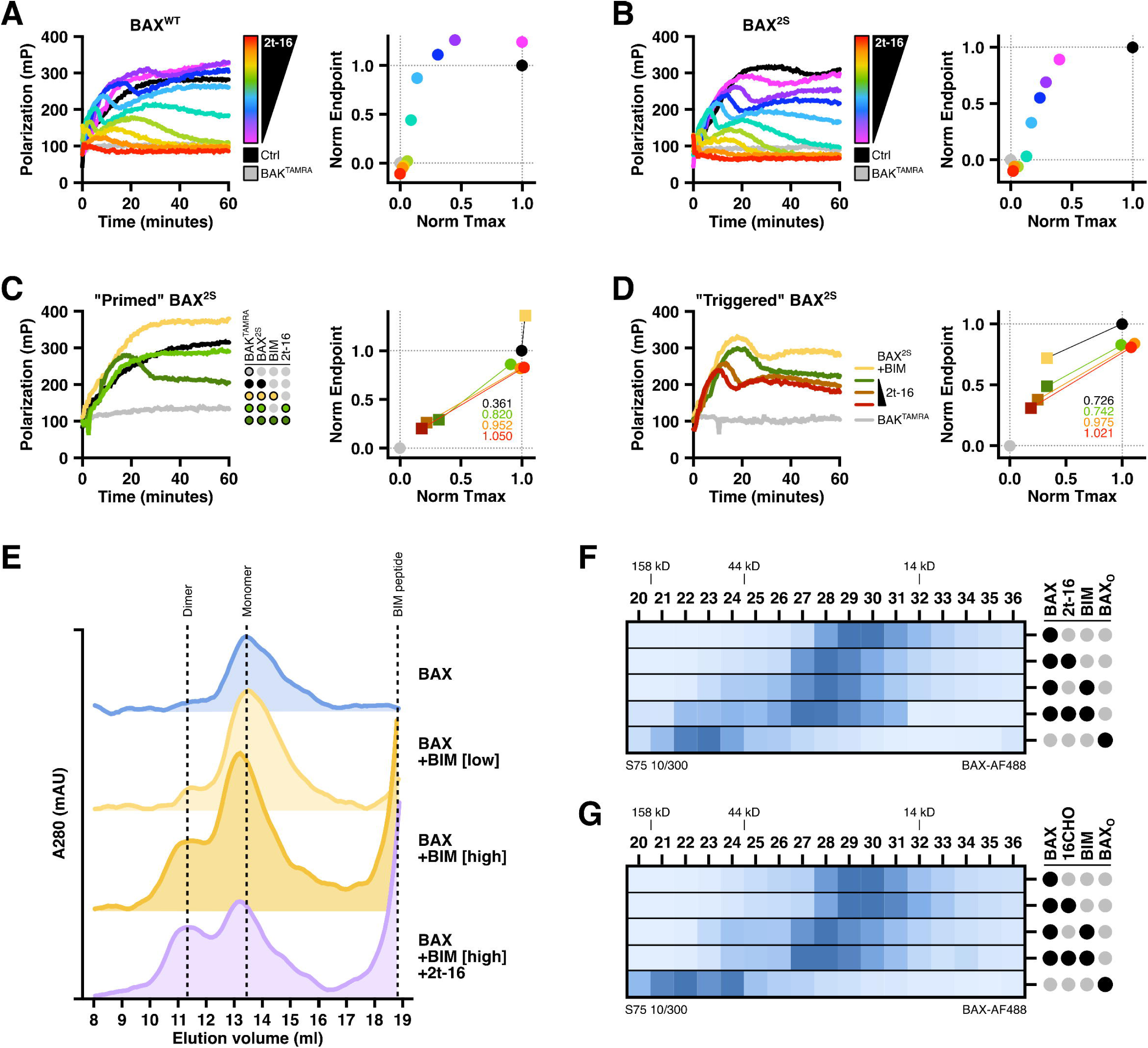
2t-hexadecenal synergizes with BIM to induce intramolecular rearrangements and early-activation of monomeric BAX. **(A–D)** BAX was treated as indicated, combined with a TAMRA-labeled BAK–BH3 peptide (BAK^TAMRA^), and immediately subjected to FLAMBE analysis in which the association and activation-induced dissociation of BAK^TAMRA^ is monitored via changes in Polarization. Left panels: kinetic Polarization data; right panels: two-dimensional plot of parameterized FLAMBE data comparing Tmax and endpoint Polarization metrics normalized to BAK^TAMRA^ (grey data) and the vehicle-treated BAX control (black data). **(A)** BAX^WT^ (60 nM) was treated with 2t–16 (2–50 μM), combined with BAK^TAMRA^ (50 nM), and subjected to FLAMBE. **(B)** Same as in **A** with BAX^2S^ (60 nM). **(C)** Left: BAX^2S^ (60 nM) was combined with non-activating concentrations of BIM–BH3 peptide (0.15 μM) and 2t-16 (4.5 μM), followed by BAK^TAMRA^ (50 nM), and subjected to FLAMBE. Right: Parameterized FLAMBE data including three concentrations of 2t–16 (green: 4.5 μM; orange: 6.5 μM; red: 10 μM) in the absence or presence of BIM–BH3 (circle and square datapoints, respectively). Annotations report the magnitude of shift between data with and without BIM–BH3. **(D)** Left: Same as in **C** with an activating concentration of BIM–BH3 and three concentrations of 2t–16 (green: 4.5 μM; orange: 6.5 μM; red: 10 μM). Right: Parameterized FLAMBE data reporting shift in samples with or without BIM–BH3. Conditions without BIM–BH3 provided in Figure **S3F**. **(E)** BAX^WT^ (800 nM) was treated with BIM–BH3 (2.5, 10 μM) and 2t–16 (50 μM) for 1 hour, subjected to size exclusion chromatography, and measured by 280 nm absorbance (A280) as eluate flowed out of the column. **(F)** Alexa Fluor 488-labeled BAX^WT^ (400 nM) was treated with 2t-16 (50 μM) and/or BIM–BH3 (10 μM) for 1 hour and subjected to size exclusion chromatography. Samples of fractions were analyzed by fluorescence spectroscopy to track BAX. Oligomeric BAX (BAX_O_) was generated via overnight treatment with DDPC (1 mM). **(G)** Same as in **F** with 16CHO (50 μM). See also Figure S3.

The consequence of BAX activation is translocation to the OMM and oligomerization, and while it is largely agreed that high-molecular weight oligomers are formed within the OMM, there is evidence supporting that physiologically-activated BAX forms low-order multimers (e.g., dimers) in solution prior to integrating with membranes.^6,8,27^ We investigated the consequence of hexadecenal-mediated synergy with BIM-activated BAX by performing size exclusion chromatography (SEC) and observed a substantial shift from monomeric to dimeric BAX when co-treated with BIM–BH3 peptide and hexadecenal (Figure 3E). To confirm that the earlier peak was BAX, we treated fluorescently-labeled BAX, subjected it to SEC, and screened the fractions for fluorescence. The monomeric BAX peak (fractions 28–31) shifted slightly left upon addition of hexadecenal or BIM–BH3 peptide (fractions 27–30), likely due to binding-dependent changes in molecular volume, and dimeric species were observed in the BIM-treated sample (fractions 23– 26); the co-treatment of hexadecenal and BIM–BH3 resulted in an increased shift and intensity indicating a greater percent of the BAX population formed multimeric species (fractions 22–26) (Figure 3F). By contrast, no such shift in the monomeric peak or BIM-induced dimer peak was observed with hexadecanal (Figure 3G). Of note, BAX activated in solution does not readily form high-molecular weight species without the stabilizing and concentrating influence of a hydrophobic environment (e.g., a membrane or micellar detergent)^8,28^, though as a reference we were able to observe BAX oligomers generated with a detergent (BAX_O_).^29^ These data collectively demonstrate that hexadecenal promotes monomeric BAX activation downstream of BCL–2 protein interactions and following activation by direct activators.

### An alpha 8 helix-gated funnel-shaped hydrophobic cavity in the BAX core interacts with 2t–hexadecenal

Hexadecenal synergized with BIM-mediated BAX activation and we observed no evidence of competition with the BAK^TAMRA^ peptide in our FLAMBE assays, suggesting that hexadecenal was binding to a site distinct from either the trigger site or BC groove, the two BH3-interacting sites respectively.^30,31^ Despite having a relatively smooth surface and no obvious “binding pocket,” studies have identified small molecules that bind to BAX, either at BH3-interacting sites or allosterically.^32–34^ To identify putative interaction sites, we performed 2D ^1^H-^15^N heteronuclear single quantum coherence (HSQC) nuclear magnetic resonance (NMR) of ^15^N-labeled BAX^WT^ treated with hexadecenal and measured a multitude of peak shifts (Figures S4A−B). Several residues exhibited significant chemical shift perturbations (CSPs) in response to hexadecenal, some of which were within unstructured regions (such as the N-terminus) or highly exposed areas (such as α4 and α9), but notably the two cysteine residues (C62 and C126) did not reach significance (Figure 4A). Interestingly, several shifts were observed in core-facing residues of α2, α5, α6, and α8, as well as bulky residues proximal to α8 within the α4–5 loop and α7. To determine whether the shifted residues were specific to the activation mechanism, we compared this CSP profile against CSPs calculated from ^1^H-^15^N HSQC NMR of BAX^WT^ treated with the non-activating saturated aldehyde. Residues exhibiting significant shifts in response to hexadecanal were more localized to accessible regions of BAX along α2, α3, and α9 (Figure S4C). Comparing the CSPs of the two aldehydes highlighted that many of the hexadecenal-induced shifts were not observed with the saturated aldehyde (Figure S4D). These core-facing residues are not believed to be readily accessible and therefore were likely to be meaningful for hexadecenal interaction and function.

**Figure 4:**
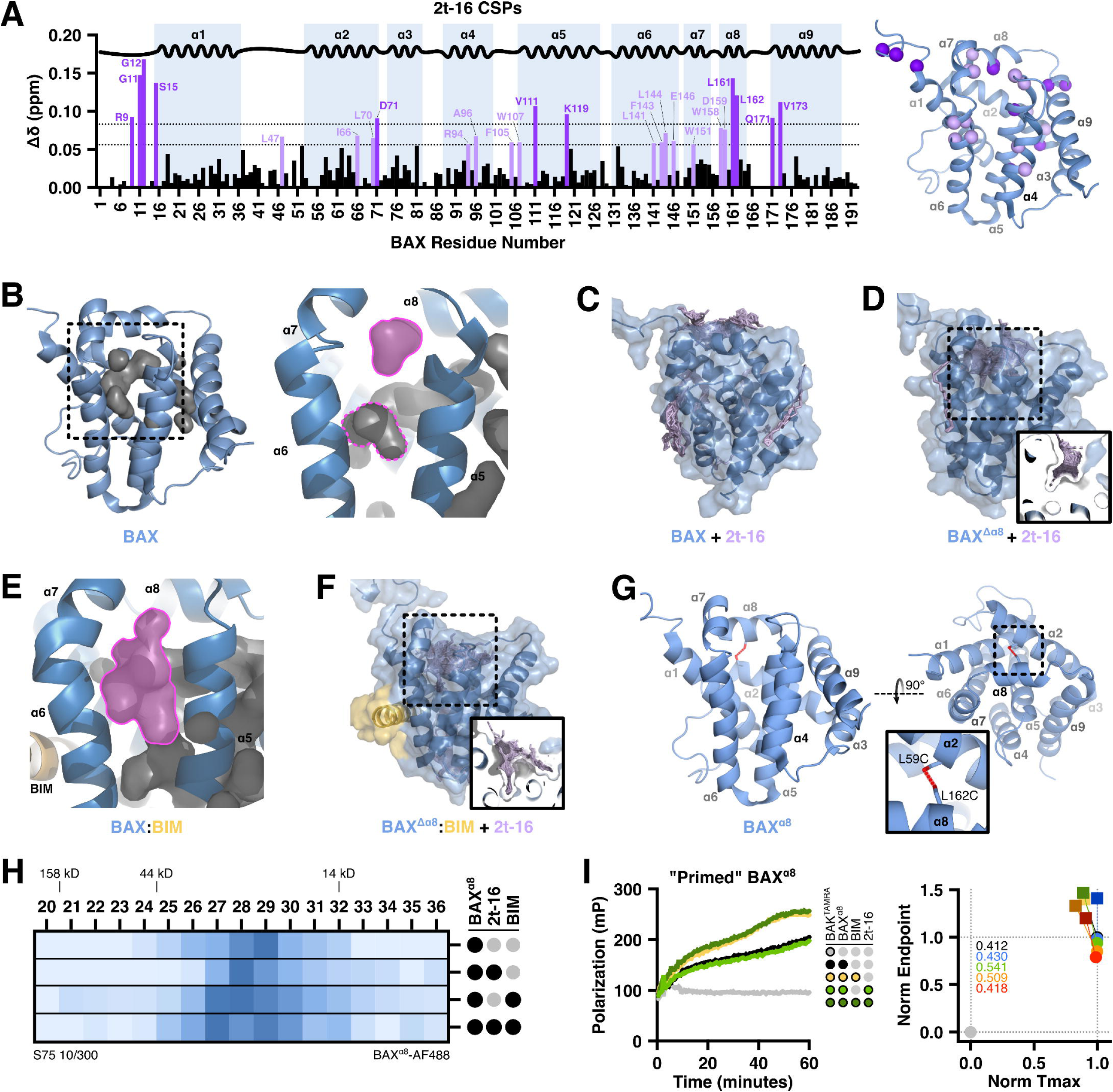
2t–hexadecenal interacts with an alpha 8 helix-gated hydrophobic cavity in the core of BAX. **(A)** ^15^N-labeled BAX^WT^ (40 μM) was treated with vehicle or 2t–16 (50, 150 μM) and subjected to ^1^H-^15^N HSQC NMR. Chemical shift perturbations (CSPs) are plotted as a function of BAX residues. Residues exhibiting a shift greater than 1 or 2 standard deviations above the average (dotted lines) are colored light and dark purple, respectively, and indicated on the BAX structure (PDB: 1F16). The absence of a bar indicates no chemical shift difference, the presence of a proline, or the residue could not be definitively assigned. **(B)** Cartoon visualization of BAX (PDB: 1F16) and the hydrophobic cavity in the BAX core formed by helices α1, α5, α6, and α7. The cavity proximal to α8 is highlighted; a distinct cavity deeper in the core between α5 and α6 is outlined. Cavity determination and visualization was performed with PyMOL using a cavity radius of 3 and a cavity cutoff of -5.5. **(C)** Unbiased *in silico* rigid docking of 2t–16 against the unmodified BAX structure (PDB: 1F16) using the SwissDock web service. Results position the majority of 2t–16 poses at the α8 helix. **(D)** *In silco* docking was performed as in **C** on a BAX structure with α8 removed (PDB: 1F16, Δ157–163; “BAX^Δα8^”). Inset displays cross-section of 2t–16 docking within the hydrophobic funnel. **(E)** Unmodified structure of BAX bound to a BIM–BH3 peptide (PDB: 2K7W; “BAX:BIM”) with the enlarged, connected cavity visualized as in **B**. **(F)** Results of 2t–16 docking on the structure of BAX:BIM with α8 removed (PDB: 2K7W, Δ157– 163; “BAX^Δα8^:BIM”) as in **C**. Inset displays cross-section of ligand docking within the hydrophobic funnel. **(G)** Cartoon visualization of the BAX^α8^ mutant (modified from PDB: 1F16). Residues L59 and L162 were mutated to cysteine and oxidized to form a disulfide tether immobilizing α8. **(H)** Alexa Fluor 488-labeled BAX^α8^ (400 nM) was treated with 2t-16 (50 μM) and/or BIM–BH3 (10 μM) for 1 hour and subjected to size exclusion chromatography. Samples of fractions were analyzed by fluorescence spectroscopy to identify BAX-containing fractions. **(I)** Left: BAX^α8^ (225 nM) was combined with non-activating concentrations of BIM–BH3 peptide (0.15 μM) and 2t-16 (4.5 μM), followed by BAK^TAMRA^ (50 nM), and subjected to FLAMBE analysis. Right: Parameterized FLAMBE data including four concentrations of 2t–16 (blue: 3 μM; green: 4.5 μM; orange: 6.5 μM; red: 10 μM) in the absence or presence of BIM–BH3 (circle and square datapoints, respectively). Annotations report the magnitude of shift between data without and with BIM–BH3. Kinetic data for all concentrations is provided in Figure **S4J**. See also Figure S4.

Inspection of the BAX structure revealed a cavity formed by hydrophobic residues in the core of the protein.^6^ The cavity shape resembles a funnel and is comprised of two topographies: a wide and shallow “mouth” formed by residues in α1, α2, α5, and α8; a narrow “neck” extending into the BAX core between α1, α5, and α6 (Figure 4B). We hypothesized that this hydrophobic “funnel” could be a desirable interaction site for hexadecenal – an inherently lipidic aldehyde – and therefore we performed unbiased *in silico* docking simulations using the SwissDock web service to model interactions with BAX. Binding modalities of hexadecenal were clustered into a few distinct regions on BAX, but 59.4% of the poses were proximal to α8, aligning with the CSPs observed by NMR (Figure 4C).

Our functional interrogations indicated that hexadecenal synergizes with BIM-mediated BAX activation, and we reasoned that BIM-induced intramolecular arrangements may induce flexibility and/or mobility of α8, making the funnel accessible to hexadecenal. SwissDock utilizes rigid receptor docking, which results in the α8 helix firmly blocking the funnel. To create a funnel-accessible structure, we removed α8 (BAX^Δα8^) and docking against the BAX^Δα8^ structure revealed a clear preference for the funnel with 97.3% of hexadecenal poses positioned within the funnel (Figure 4D). Interestingly, when we examined the solution NMR structure of BAX bound to a stapled BIM–BH3^30^, we observed an enlargement of the hydrophobic funnel, most notably in the neck of the funnel (Figure 4E). In the unbound BAX structure, the neck of the funnel is cinched into discontinuous cavities by residues of α5 and α6; in contrast, the BIM-bound structure has a connected cavity due to a shift in the residues lining the funnel, increasing both the depth and width. Docking hexadecenal against the BAX:BIM structure with α8 removed (BAX^Δα8^:BIM) revealed that the aldehyde was positioned in the funnel, frequently posed as being inserted into the neck of the funnel (Figure 4F).

To substantiate our *in silico* conclusions, we engineered a structural mutant of BAX to restrict the mobility of α8 and the subsequent access to the hydrophobic funnel. We introduced two cysteine residues into BAX^2S^ (L59C, C62S, C126S, L162C; “BAX^α8^”) that could be oxidized to induce a disulfide tether between α2 and α8 (Figure 4G). We characterized the consequence of the new mutations by activating BAX^α8^ with BIM–BH3 peptide and assessing LUV permeabilization. Reduced BAX^α8^ (i.e., “unlocked”) remained functional and exhibited BIM-induced LUV permeabilization (Figure S4E); in contrast, oxidized BAX^α8^ (i.e., “locked”) demonstrated no permeabilization (Figure S4F). Several studies have characterized that BAX functionalization and pore formation required large-scale conformational reorganization, which involves the separation of α8 from core helices, and therefore it is not surprising that BAX^α8^ was a “functionally dead” structural mutant.^5–7,35^ However, we recently demonstrated that several historically “dead” BAX mutants remain sensitive to BH3-induced activation and demonstrate early-activation structural hallmarks despite not maturing into functional pore-forming conformations.^8^ SEC experiments revealed that locked BAX^α8^ exhibited a shift in response to BIM–BH3 but was unaffected by either hexadecenal alone or coupled with BIM-mediated activation (Figure 4H). To further investigate whether the immobilized α8 helix would disrupt sensitivity to hexadecenal, we studied BAX^α8^ activation using our FLAMBE assay. Both the unlocked and locked forms of BAX^α8^ demonstrated activation by BIM–BH3, indicating that that the mutant still exhibits activation-induced intramolecular rearrangements (Figures S4G–H). Critically, locked BAX^α8^ did not activate when treated with hexadecenal (Figure S4I). Moreover, BIM-primed BAX^α8^ displayed no activation or synergy in the presence of hexadecenal (Figures 4I, S4J). These data substantiate the docking simulations and the conclusion that hexadecenal-induced BAX activation is mediated through non-covalent interactions with the hydrophobic funnel in the BAX core, which is exposed following BIM triggering. Given that this hydrophobic cavity potentiates BAX activation, we termed this site the “BAX Actuating Funnel”, or BAF.

### Aldehyde length and BAF topography are determinants of BAX activation

The saturated aldehyde hexadecanal did not activate BAX suggesting that the mechanism is unique to 2t–hexadecenal. We next considered the relevance of aldehyde length on interactions with the BAF by utilizing a panel of 2t–alkenals ranging from 5 to 13 carbon chains (henceforth, “alkenals”). Experiments with a mid- or short-chain alkenal, nonenal (“2t–9”) and pentenal (“2t–5”), respectively, demonstrated CSP profiles that were similar to hexadecenal, but were also more diffuse across BAX (Figure 5A). While there was some conservation of CSPs observed in residues within and proximal to α8, the shorter alkenals exhibited a plethora of shifts within α4 and α9 as well as a general trend of interacting with solvent-exposed residues (Figure S5A). These data may also indicate that shorter alkenals are more promiscuous, as nonenal and especially pentenal interacted with several accessible regions of BAX, exhibited decreasing specificity at higher concentrations, and displayed weaker CSPs (Figures S5B–C). Docking simulations with alkenals ranging from 5 to 15 carbon chains supported this hypothesis by demonstrating a length-dependent specificity for the α8 region (in the full-length structure, BAX) or the BAF (in the BAX^Δα8^ structure) (Figures 5B, S6A–B). When docked against the BAX^Δα8^:BIM structure, most alkenal species were predicted to interact with the larger BAF, though many were not positioned in the depth of the funnel and short-chain aldehydes could not occupy both the mouth and neck of the BAF in the same pose (Figure S6C). In the BIM-bound BAX structure, mobilization of the α1–α2 loop created a region that was predicted to be an interaction site, and longer aldehyde structures were disproportionately positioned as a consequence; we believe this was an artifact of rigid receptor docking and not a physiological phenomenon, and therefore we did not quantify the percentage of binding poses localized to the BAF. The precursor of 2t–hexadecenal, sphingosine 1-phosphate (S1P), was previously identified as a requirement for BAK-mediated MOMP but did not directly promote BAX-mediated MOMP.^15^ We docked the structure of S1P against BAX models and observed a reduced preference for the BAF and the tail could not occupy the neck of the BAF when modeled against BAX^Δα8^:BIM structure, likely due to the bulky and negatively charged phosphate group (Figures 5B, S6A–C). Collectively, NMR and unbiased docking assessments supported our hypothesis that aldehyde length promotes specificity for the BAF and BAX activation.

**Figure 5:**
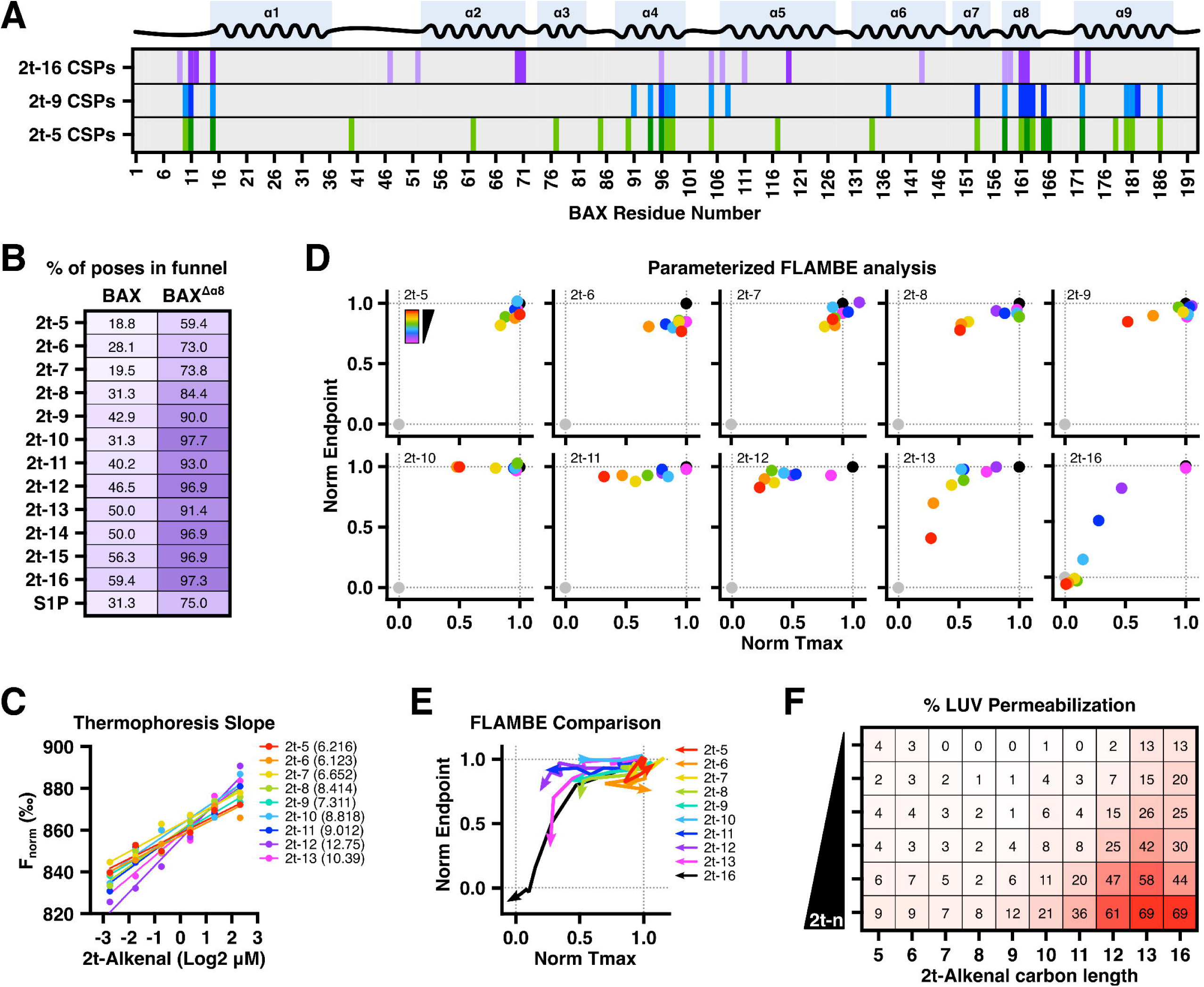
Aldehyde length controls BAF function and BAX activation. **(A)** ^15^N-labeled BAX^WT^ (40 μM) treated with 2t–16, nonenal (2t–9), or pentenal (2t–5) and subjected to ^1^H-^15^N HSQC NMR. Residues exhibiting significant CSPs averaged across concentrations for each of the indicated 2t–alkenal are highlighted. Each row is color-coded to report residues exhibiting shifts greater than 1 (light color) or 2 (dark color) standard deviations above the average across measurable shifts. Data summarized from Figure **S5A**. **(B)** Results of unbiased *in silico* rigid docking 2t–alkenals or S1P against the full-length BAX or BAX^Δα8^ structure using the SwissDock web service. Binding modalities were inspected for positioning proximal (for BAX^FL^) or within (BAX^Δα8^) the BAF and reported as the percent of total poses. Visualizations of docking results provided in Figure **S6A–B**. **(C)** Summary of MST analyses with BAX^WT^ incubated with 2t–alkenals of differing carbon lengths (5–13 carbons) at a range of concentrations (0.16–5 μM). The thermophoresis datapoints were fit and the slope is provided for each alkenal. Original timetrace data provided in Figure **S7A**. **(D)** Parameterized data from FLAMBE studies with BAX^2S^ (60 nM) treated with 2t–alkenals of differing carbon lengths (5–13, 16 carbons) at several concentrations (3–50 μM). Black and grey datapoints report vehicle-treated BAX^2S^ and BAK^TAMRA^, respectively. Original data provided in Figure **S7B**. **(E)** FLAMBE data trends for each of the 2t–alkenal titration from **D**. **(F)** Heatmap visualization of endpoint permeabilization data from LUVs incubated with BAX^2S^ (100 nM) treated with 2t-alkenals of differing carbon lengths (5–13, 16 carbons) at several concentrations (6.5–50 μM). Each experiment was normalized to the matching vehicle-treated BAX^2S^ condition to control for variability. Original data provided in Figure **S7C**. See also Figures S5−7.

Next, we tested our hypothesis by investigating which, if any, alkenals could phenocopy hexadecenal function and BAX activation. When subjected to MST, BAX^WT^ exhibited altered thermophoresis with each of the alkenals; though the fit of the data demonstrated a trend in which the slope increased with aldehyde length (Figures 5B, S7A). To interrogate the functional consequence of alkenals interacting with BAX, we next subjected each alkenal to FLAMBE analysis and observed a dependency on aldehyde length for BAX^2S^ activation (Figure 5D). Interestingly, at the concentrations we tested, undecenal (“2t–11”) and dodecenal (“2t–12”) displayed some disruption of BAX^2S^:BAK^TAMRA^ interactions, similar to the results with hexadecanal, but BAK^TAMRA^ was able to re-bind and did not exhibit an activation profile (Figures S3E, S7B). When compared to hexadecenal, only tridecenal (“2t–13”) exhibited a clear dose-dependent activation trend, albeit less potently than hexadecenal (Figures 5D–E, S7B). To definitively determine alkenal-induced BAX activation, we performed LUV permeabilization studies and confirmed a length-dependent sensitivity for BAX^2S^-mediated pore formation (Figures 5F, S7C). Additionally, these results were conserved in experiments using BAX^WT^ (data not shown). Taken together with our structural and modeling data, these results indicate that chain length is a determinant of α,β-unsaturated alkenals to specifically target the BAF and activate.

Having identified determinants of ligand specificity, we next sought to characterize the residues of the BAF that regulate interactions with, and sensitivity to, hexadecenal. We performed molecular docking of hexadecenal against the BAX:BIM structure using the GLIDE software with a docking grid centered on the BAF. As we expected, the residues lining the BAF that interact with hexadecenal were predominantly hydrophobic residues of α5 and α6 and all predicted poses were positioned into the neck of the BAF (Figures 6A−B). We performed a virtual mutagenesis screen using the BAX structure and identified that mutating V110, L113, or L144 to bulkier residues was predicted to alter the BAF (Figure 6C). Indeed, performing molecular docking on these mutant structures demonstrated worse binding scores and altered hexadecenal positioning due to the disruption of BAF topography (Figure 6D−E). Of note, we ran clustering analysis on all the docked poses for each mutant and found that the poses were sufficiently similar to be clustered together with the exception of BAX^L113M^, which still positioned one pose into the shallower BAF; however, this pose had worse scoring compared to the top L113M hits.

**Figure 6:**
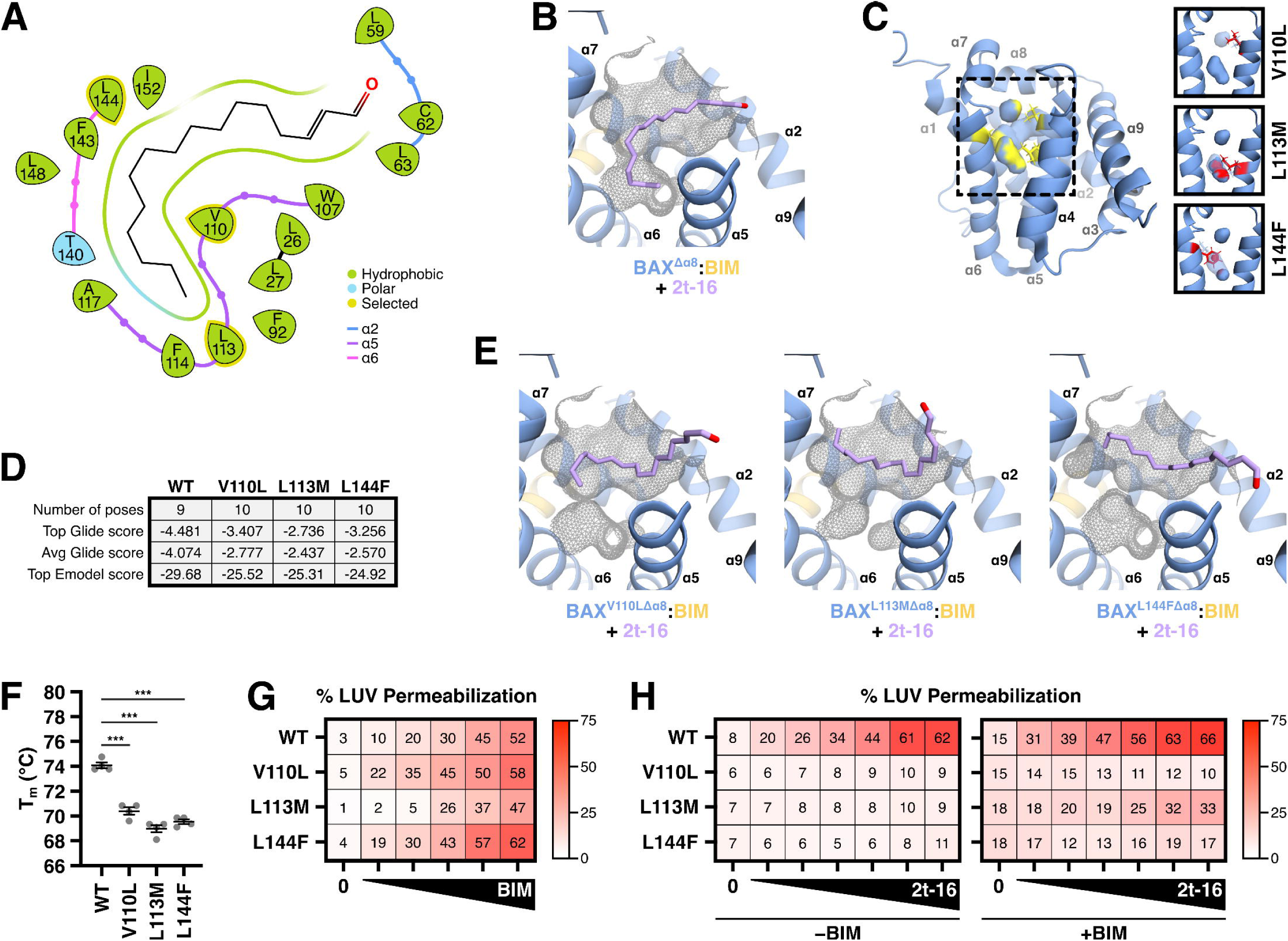
Proximal BAF mutations disrupt 2t–hexadecenal synergy with BAX and BIM. **(A)** Molecular docking and modeling of BAX^Δα8^:BIM (PDB: 2K7W, Δ157–163) with 2t–16 using Schrödinger Glide with a binding region localized to the BAF. The interaction diagram identified several hydrophobic residues lining the BAF. **(B)** The highest scoring pose from **A** is shown and is representative of all generated poses. Surface of exposed BAF shown in grey wireframe and clipped to aid in visualization. **(C)** Cartoon visualization of full-length BAX (PDB: 1F16) with BAF residues identified in **A** highlighted in yellow. A virtual mutagenesis screen was performed with PyMOL to identify BAF-disrupting mutations. Right: candidate side chain replacements altering the BAF are shown in red and compared to the WT BAF. Cavity determination and visualization was performed with PyMOL using a cavity radius of 3 and a cavity cutoff of -5.5. **(D)** Molecular docking as in **A** against structures containing the indicated mutations. Generated poses of 2t–16 had reduced interaction scores against mutant structures compared to the wild type structure resulting from the altered BAF. **(E)** The highest scoring poses from **D** are shown and representative of generated poses. **(F)** The melting temperature of each mutant was measured by thermal shift differential scanning fluorimetry using SYPRO orange. Statistical significance was determined by two-way ANOVA; *** denotes *P* < 0.001. **(G)** Heatmap visualization of normalized endpoint LUV permeabilization data for WT and BAF mutants activated with a range of BIM–BH3 concentrations (0.125−2 μM). Data summarized from Figure **S8A**. **(H)** Heatmap visualization of normalized endpoint LUV permeabilization data from LUVs incubated with the BAF mutants treated with 2t–16 (6.5–50 μM) ± BIM–BH3 peptide (0.15 μM). Data summarized from Figure **S8B−C**. See also Figure S8.

To corroborate our *in silico* predictions, we generated recombinant protein for these BAF mutants and tested them for BAX functionality. Several established BAX point mutants result in a deficient or functionally dead protein, and so we first determined the melting temperature of our BAF mutants. Compared to WT, each BAF mutant exhibited a lower melting temperature, suggesting that these mutations would not be functionally deficient (Figure 6F). While the lower melting temperature may also suggest reduced stability of the monomeric protein, we did not observe any redistribution to oligomeric species during protein purification and storage (data not shown). Furthermore, we confirmed that each BAF mutant responded to BIM activation with similar kinetics and endpoint, noting only slight insensitivity of BAX^L113M^ at low BIM concentrations but similar activation at high concentrations (Figures 6G, S8A). Critically, each BAF mutant demonstrated complete insensitivity to hexadecenal and exhibited no synergy with BIM-primed BAX (Figures 6H, S8B−C). Indeed, this result was conserved even with activating concentrations of BIM (data not shown). It is worth noting that BAX^L113M^ did retain some synergy of BIM and hexadecenal, albeit dramatically reduced, and we suggest that this may be a consequence of a shallower, but relatively intact BAF in the BAX^L113M^ structure (Figure 6E). As an aside, we did generate and screen candidate mutations altering the bulky BAF residues (i.e., F114 and F143), but observed reduced stability, oligomerization during purification, and substantially increased sensitivity to BIM, which obfuscated interpretations with hexadecenal (data not shown). Interestingly, a prior study also noted that mutation of F114 is highly sensitizing,^36^ and we posit that substituting these bulky residues enlarges the BAF and subsequently destabilizes BAX (see Discussion). Collectively, these investigations indicate that the activating mechanism of hexadecenal is the ability of the aldehyde to reside in the depth of the BAF, and alteration to the BAF shape or aldehyde size disrupts this fundamental interaction.

### Proline 168 allosterically controls the BAF and 2t–hexadecenal function

Our modern understanding of the BAX structure-function relationship includes the concept of allostery, in which interactions on BAX can cause structural rearrangements in distal regions. This concept is observed during BIM-mediated activation, which elicits mobilization of α9 from the BC groove^30^, or binding of a sensitizing molecule to the hairpin pocket, which provokes exposure of the α1−2 loop^33^ – both structural events are hallmarks of BAX activation.^4^ Recently, the structure of BAX containing a mutation of proline 168 (BAX^P168G^) was described and proposed to cause rotation of bulky sidechains in the allosteric α4−5 and α7−8 loops.^37^ Despite identification of P168 as critical for BAX activation, translocation, and pore-forming ability over 20 years ago,^38^ and its recent identification as an arising loss-of-function mutation conferring resistance to venetoclax therapy in acute myeloid leukemia,^39^ a mechanistic explanation for its requirement has remained elusive. We hypothesized that the alternative rotamers in bulky α8-proximal residues may consequently alter the BAF and, indeed, the neck of the BAF was lost in the BAX^P168G^ structure (Figure 7A). Furthermore, molecular docking against the BAX^P168G^ structure positioned the aldehyde with the carbon chain now pivoting towards α1 residues (Figures 7A−B). Directly comparison between poses of hexadecenal docked to WT and P168G structures revealed distinct interaction signatures favoring interactions with α5 and α6 or α1 residues, respectively (Figure 7C).

**Figure 7:**
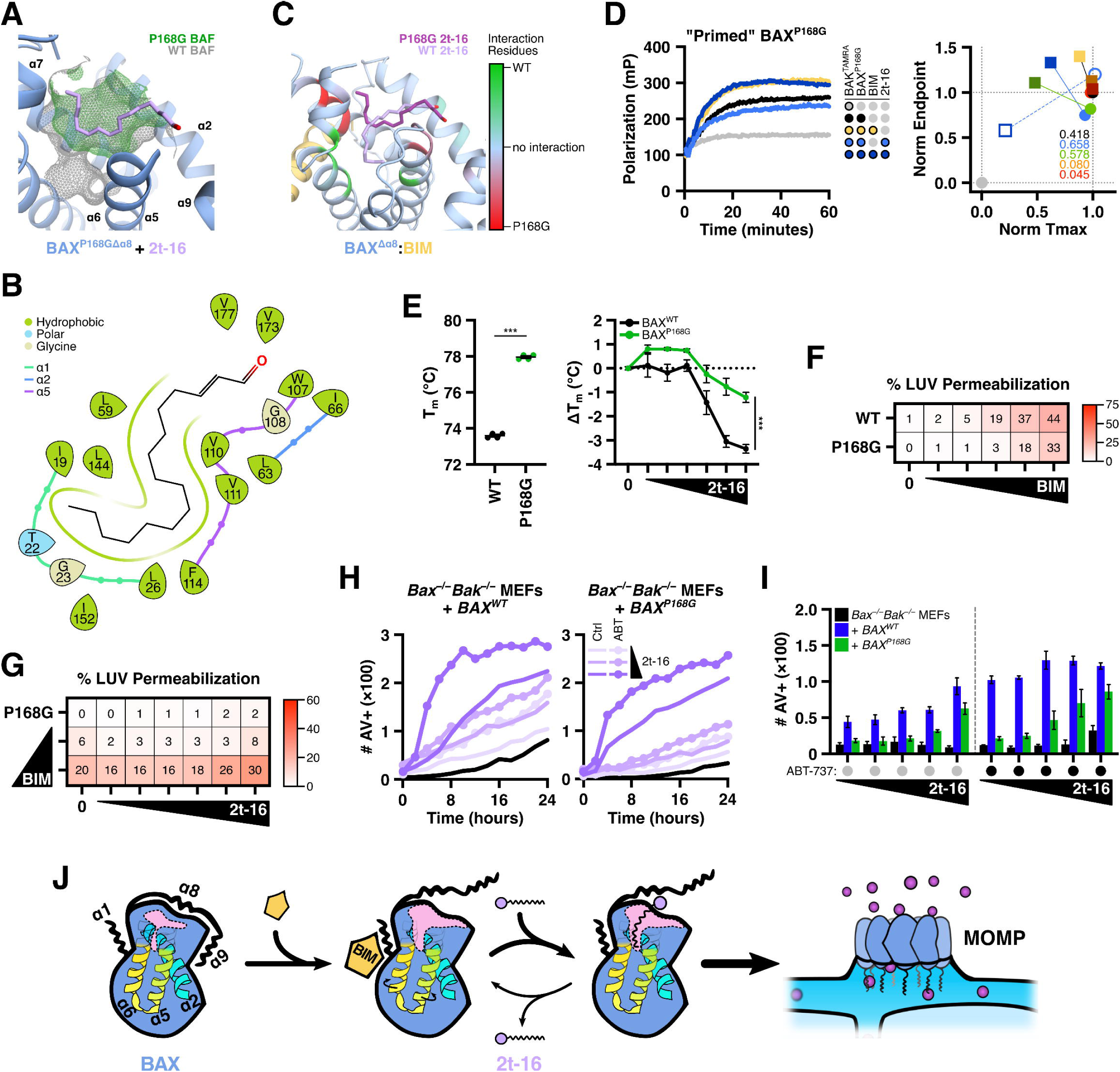
Mutation of proline 168 allosterically disrupts the BAF and 2t–hexadecenal function. **(A−C)** Molecular docking and modeling of BAX^P168^^GΔα8^ (PDB: 5W60, Δ157–163) with 2t–16 using Schrödinger Glide with a binding region localized to the BAF. **(A)** The BAX^P168G^ structure includes divergent rotamers of bulky core residues that removes the neck of the BAF. The highest scoring pose is shown and is representative of generated poses. Surface of exposed BAF shown in green wireframe and clipped to aid in visualization. The BAF surface from WT BAX is overlayed in grey for comparison. **(B)** The interaction diagram from **A** reveals that 2t–16 interacts with distinct residues, including sidechains of α1. **(C)** Residues interacting with 2t–16 were compared between WT and P168G BAX and colored green or red, respectively, according to the number of poses and specificity for one of the isoforms. Structure of BAX^Δα8^:BIM included for visualization as well as top scoring 2t–16 poses. **(D)** Left: BAX^P168G^ (65 nM) was combined with non-activating concentrations of BIM–BH3 peptide (0.15 μM) and 2t-16 (3 μM), followed by BAK^TAMRA^ (50 nM), and subjected to FLAMBE. Right: Parameterized FLAMBE data including four concentrations of 2t–16 (blue: 3 μM; green: 4.5 μM; orange: 6.5 μM; red: 10 μM) in the absence or presence of BIM–BH3 (circle and square datapoints, respectively). Annotations report the magnitude of shift between data with and without BIM–BH3. Empty symbols denote parameterized data for BAX^WT^ shown in Figure **S9A** and is included for comparison. **(E)** The melting temperature of BAX^WT^ and BAX^P168G^ ± 2t–16 (6.5−50 μM) was measured by thermal shift assay using SYPRO orange and compared. Statistical significance was determined by two-way ANOVA; *** denotes *P* < 0.001. **(F)** Heatmap visualization of normalized endpoint LUV permeabilization data for BAX^WT^ and BAX^P168G^ mutants activated with a range of BIM–BH3 concentrations (0.125−2 μM). Data summarized from Figure **S9B**. **(G)** Heatmap visualization of normalized endpoint LUV permeabilization data from LUVs incubated with the BAF mutants treated with 2t–16 (6.5–50 μM) ± BIM–BH3 peptide (0.15 μM). Data summarized from Figure **S9C**. **(H−I)** *Bax^−/−^Bak^−/−^* double knockout MEFs reconstituted to express BAX^WT^ or BAX^P168G^ were subjected to SPARKL analysis measuring real-time labeling with fluorescently-tagged Annexin V (100 μg/ml) via imaging with a Cytation 7. Data are reported as positive events per image. **(H)** Kinetics of cell death for reconstituted MEFs treated with 2t–16 (10−30 μM) and co-treated with DMSO vehicle or ABT-737 (1 μM); black lines report vehicle control. Data are the mean of replicates and error bars are omitted for visualization. **(I)** Comparison of Annexin V labeling at 18 hours for parental *Bax^−/−^Bak^−/−^* MEFs and BAX^WT^ or BAX^P168G^ reconstituted MEFs. Data shown are the mean of technical replicates and error bars report SEM. **(J)** A cohesive model of protein and lipid contributions to BAX activation. A cross section of BAX is illustrated to visualize changes to the hydrophobic funnel shape and accessibility (highlighted in pink). See also Figure S9.

We reasoned that disruption of the BAF in BAX^P168G^ would inhibit interactions with hexadecenal. Using our FLAMBE assay, we demonstrated that BIM-primed BAX^P168G^ did not activate in response to hexadecenal as compared to BAX^WT^ (Figures 7D, S9A). Furthermore, BAX^P168G^ exhibited less shift in melting temperature due to hexadecenal (Figure 7E). As expected, BAX^P168G^ was less sensitive to BIM activation and resulted in attenuated membrane permeabilization, but notably was not entirely functionally dead (Figures 7F, S9B). In contrast, BAX^P168G^ demonstrated no activation in response to hexadecenal and only minor synergy with BIM at activating concentrations, suggesting that activation-induced molecular rearrangements are the limiting factor for the synergistic function of hexadecenal (Figures 7G, S9C). As detailed by others,^37^ we also observed a significantly higher melting temperature in BAX^P168G^ compared to BAX^WT^, suggesting a stabilization of the monomeric conformer (Figure 7E). Finally, we replicated our apoptosis experiments in *Bax^−/−^Bak^−/−^* DKO MEFs transduced to stably express either BAX^WT^ or BAX^P168G^ (Figure S9D). The DKO MEFs reconstituted with BAX^WT^ exhibited greater sensitivity to ectopic hexadecenal compared to BAX^P168G^-expressing MEFs (Figure 7H). Furthermore, this discrepancy was maintained in MEFs co-treated with hexadecenal and ABT–737 (Figures 7H−I). Taken together, these results reveal that BAX^P168G^ is insensitive to hexadecenal and synergy with BIM due to disfunction of the BAF. Furthermore, we posit that the increased flexibility in the α8−9 loop (by substituting a rigid proline for glycine) may also alter the mobilization of α8 and subsequent BAF exposure following BIM triggering in addition to deforming the BAF structure through allosteric sidechain reorganization.

## DISCUSSION

### 2t–hexadecenal non-covalently binds the BAX Actuating Funnel to potentiate BAX activation

Several decades have been devoted to characterizing the protein-protein interactions that govern the BCL-2 family and mediate the onset of MOMP following pro-apoptotic signaling. More recently, a growing focus on the mitochondrial contributions to potentiate MOMP have been identified, including mitochondrial mass, shape or curvature, OMM-IMM junctions, mitochondrial-ER contact sites, and the OMM lipid milieu and cardiolipin.^40^ Additionally, the sphingolipid metabolites S1P and 2t–hexadecenal were identified as the first *de facto* signaling lipid species to modulate the BCL–2 effector proteins, BAK and BAX, respectively.^15^ Here, we have demonstrated that 2t–hexadecenal directly promotes BAX activation via non-covalent interactions (Figures 1–2), and cooperates with direct activators to enhance early-activation steps of monomeric BAX prior to membrane interactions or oligomerization (Figure 3). We determined that 2t–hexadecenal interacts within a hydrophobic funnel-shaped cavity in the core of BAX, formed residues of α1, α5, α6, and capped by α8, which we now term the “BAX Actuating Funnel” or BAF, and that this region is exposed and enlarged following BIM triggering (Figure 4). We identified chemical and structural determinants of the hexadecenal:BAF interaction, including aldehyde length (Figure 5) and BAF residues (Figure 6), and further revealed that proline 168 allosterically regulates the BAF and subsequent mutation reduces BAF topography and availability (Figure 7). We therefore propose a cohesive model for successive BAX activation: BIM-mediated triggering induces α9 mobilization as well as tandem α8 movement, resulting in accessibility of the BAF, interactions with 2t–hexadecenal, and promoting subsequent conformational changes and pore formation (Figure 7J).

In contrast, two previous studies reported that 2t–hexadecenal covalently modifies BAX cysteine residues through Michael addition. One study synthesized a clickable alkyene analogue of 2t–hexadecenal and observed modification of C62 in cell lysates and recombinant protein^26^, while another study observed alkylation at C126 using recombinant BAX^23^. Both studies also demonstrated that the saturated aldehyde hexadecanal was non-activating and did not modify BAX. One possible explanation is the concentration of 2t–hexadecenal used, which often ranged from high micromolar to millimolar ranges and liposome formulations upwards of 10% 2t–hexadecenal, which may have facilitated covalent modification; another is that cell lysate and recombinant protein buffers contained NP-40 nonionic detergent, which directly engages monomeric BAX activation.^2,41^ We also cannot rule out the possibility of additional differences in solvents, buffer formulations, reagents, or pH that likely affect the reaction chemistry and may explain the divergent results; however, we were unable to identify a set of conditions to induce modification by 2t–hexadecenal as a positive control for LC-MS analysis. Another difference between studies was the BAX protein, which was either tagged (HA-tagged for overexpression or His-tagged for recombinant vectors)^26^ or was determined to have a molecular weight slightly greater than the calculated mass of BAX (21320 Da observed compared to 21184 Da predicted)^23^. Perhaps these minor modifications to the BAX protein altered cysteine exposure, the neighboring residues, or otherwise favored the covalent reaction mechanism.

Critically, we do not dispute the findings of these prior studies nor the conclusion that cysteine modification modulates BAX activity.^42^ There is substantial evidence that 2t–hexadecenal and α,β-unsaturated aldehydes are inherently reactive molecular species that form adducts with a variety of macromolecules.^22,25^ In fact, over 500 proteins were identified to be modified by the 2t–hexadecenal analogue probe^26^, suggesting that cysteine modification of BAX is not a unique or selective phenomenon, which further highlights the need to individually investigate each protein. In a cell, 2t–hexadecenal is promptly metabolized by FALDH/ALDH3A2 to avoid accumulation of reactive fatty aldehydes, lipid peroxides, and alkylated adducts^16,43,44^; as such, cellular BAX is unlikely to experience micro- or millimolar concentrations of 2t–hexadecenal and subsequent modification under physiological conditions. We posit that the confluence of sphingolipid metabolism, lipid transfer, and BAX apoptotic foci forming at MERCS serves to regulate the availability of 2t–hexadecenal during BAX activation.^40^ We also observed that short-chain alkenals did not activate BAX, which aligns with a prior study that also demonstrated increased reactivity with short-chain alkenals.^23^ While not measured for 2t–hexadecenal or long-chain alkenals specifically, it has been reported that 2t–alkenals exhibit reduced reactivity in a length-dependent manner despite having similar electrophilic indices, possibly due to differences in steric hinderance or relative solubility/hydrophobicity.^45,46^ Therefore, we propose that while 2t–hexadecenal can covalently modify BAX *in vitro* and cell lysates, this likely does not represent the primary mechanism promoting BAX activation, which we suggest is via non-covalent interactions within the BAF.

### 2t–hexadecenal and the gatekeeper α8 helix in BAX activation

So how does 2t–hexadecenal binding to the BAF promote BAX activation? One possible explanation is that residence of 2t–hexadecenal in the BAF occupies space that would otherwise accommodate bulky aromatic residues of α8. Activated BAX monomers can be “reset” by anti-apoptotic BCL–2 protein (e.g., BCL–xL)^47,48^ and perhaps the steric interference of 2t–hexadecenal prevents replacement of α8 and α9 into the BAF and BC groove, respectively. Another explanation could be that binding of 2t–hexadecenal, particularly in the neck of the BAF, which resides deep in the core of BAX, is inherently destabilizing to the packing of side chains and core helices. Therefore, 2t–hexadecenal binding may aid separation of the core and latch domains and facilitate forming multimeric conformers. Additionally, residence in the BAF may also aid BAX activation by promoting exposure of the BAX BH3 residues, which are core-facing in the inactive conformer, through steric interference with α2.^4,49,50^

Interestingly, the concept of a lipid-interacting funnel cavity capped by a gatekeeper helix is also present in the only other protein known to non-covalently bind 2t–hexadecenal – FALDH. In FALDH, the substrate site is a deep hydrophobic funnel capped with a short gatekeeping helix, which is hypothesized to participate in substrate specificity.^51^ The gatekeeper helix has several aromatic residues and is stabilized through interactions with the preceding helix, which has a perpendicular orientation, and a lateral loop region, in a parallel orientation (similar to BAX α7 and α4–5 loop). Like BAX, the FALDH gatekeeper helix is the penultimate helix, and is followed by a short linker and a C-terminal helix that serves as a transmembrane domain, orienting the entrance of the funnel towards the residential membrane. Of note, FALDH functions as a symmetrical homodimer with the stabilizing helix, gatekeeping helix, and transmembrane helix folding on the trans subunit, bearing a striking similarity to the domain-swapped BAX dimer structure.^6^ The specificity of FALDH is attributed to the cooperation of the hydrophobic loop, the stabilizing helix, and the gatekeeper helix that collectively surround the substrate funnel; and it is tempting to speculate that the similar features in the BAX structure may likewise play a role in 2t–hexadecenal recruitment and specificity.

### The BAF provides structure-function insights into monomeric BAX biology

Monomeric BAX conformers must undergo substantial unfolding to adopt multimeric structures, and therefore, from a structural perspective, “activating” BAX must sufficiently destabilize the monomer to enable refolding into domain-swapped conformations. Interestingly, several features of the BAX structure support this conceptualization of activation. For example, BAX has no disulfide bonds and evolved to fold entirely through hydrophobic/hydrophilic and electrostatic interactions, which decreases the energy “hill” to transition between stable conformers (e.g., monomer to domain-swapped dimer). Additionally, cavities or voids destabilize globular proteins^52,53^, and BAX studies mutating bulky core residues with smaller side chains have observed increased sensitivity to activation^36^, likely due to enlarging the BAF and reducing stability^54,55^. Indeed, we observed this phenomenon when we mutated F114 or F143 (data not shown). Of note, the cavity has also been observed in the BID–BH3:BAX^ΔC21^ and BAX–BH3:BAX^ΔC21^ domain-swapped dimer structures (PDBs 4BD2 and 4BD6, respectively)^6^, as well as the BH3-in-groove:BAX^ΔC26^ complexes (PDBs 4ZIF and 4ZIG)^56^; albeit the location is more central to α2, α5, and α8 (i.e., the “mouth” of the BAF), and may be expanded by the removal of α9 and adjacent residues. One interesting insight is that BAX^ΔC28^ exhibited a diminished cavity due to a shift in the α2 helix^56^, suggesting that bulky residues within and proximal to α8 (e.g., F165 and W170) may aid in maintaining the cavity through interactions with α2. Additionally, we explored the role of P168 in BAF shape and availability in this work. The presence of the cavity in several solved apo and BH3-bound BAX structures strongly suggests its importance to the structure-function relationship of BAX and supports the destabilization concept of BAX activation.

Our model suggests that activation-induced mobilization of α9 also results in displacement of α8, which in turn exposes the BAF and hydrophobic core. Therefore, we propose that α8 functions as a gatekeeper helix – capping the cavity, limiting solvent exposure of hydrophobic residues, and ultimately stabilizing the monomeric conformation of BAX through interactions within the helical bundle. This model is supported by the fact that the P168G mutation, which increases flexibility in the α8−9 loop and we propose limits α8 mobility, exhibits increased protein stability as the inactive, globular monomer. Furthermore, several studies have demonstrated that disruption of α8 is deleterious to monomeric BAX stability: a proline substitution in α8 (L161P) caused oligomerization and cell death^57^; removal of α8 and α9 (BAX^ΔC35^) resulted in oligomerization^58^; C-terminal truncations (BAX^ΔC21^, BAX^ΔC25^, BAX^ΔC28^) remained monomeric but deletion of Y164 (BAX^ΔC29^) caused oligomerization^6^. Furthermore, one of our BAX^α8^-locked mutagenesis strategies (BAX^W107C,Y164C^) resulted in recombinant protein purifying primarily as oligomeric species (data not shown). Notably, W107 and Y164 are predicted to exhibit pi-stacking as are several aromatic residues localized to the α4–5 loop and α7/α8. Interestingly, these segments of BAX are the BH1 and BH2 domains, suggesting that their conservation within multidomain BCL–2 family proteins is due, at least in part, to their role in stabilizing the helical bundle via the α8 cap.

### Concluding thoughts

Here, we describe a cohesive model of BAX activation, in which successive contributions of protein-protein and protein-lipid interactions promote intramolecular maturation of BAX monomers into functional oligomers. We propose that interactions between 2t–hexadecenal and the exposed BAF can occur either in the cytosol, as we have modeled herein, or at the OMM, as modeled by others^23^. Lipids play a critical role in BAX activation, both in early-activation steps as well as pore formation. Interestingly, while activated BAK structures bind and incorporate lipid moieties, analogous studies have not identified a similar mechanism in BAX multimers suggesting that there may be a divergent mechanism.^59,60^ BAX is unique in that much of its regulation occurs in solution prior to membrane association and therefore the lipid contributions to BAX activation may function primarily on monomeric activation instead of oligomerization. Finally, this work describes the function and importance of the BAF, which had previously been identified but its significance was never determined.^6,56^ The BAF represents a new regulatory region that is critical for understanding monomeric BAX structure-function biology as well as providing a new space for pharmacologically targeting BAX. The BAF is exposed following BIM- or BID-induced BAX activation and thus we suggest that BAF-targeting molecules would be particularly adept at modulating BAX in cellular contexts that actively maintain direct activator BH3–only protein function (e.g., “primed” cells) while being well tolerated in healthy tissues where the BAF is unexposed.

## ACKNOWLEDGEMENTS

This work was supported by NIH grants F31AA024681 (J.D.G.), R01-GM083159 (R.W.K.), R01-CA237264 (J.E.C.), R01-CA267696 (J.E.C.), and R01-CA271346 (J.E.C.); a Collaborative Pilot Award from the Melanoma Research Alliance (J.E.C.); a Department of Defense − Congressionally Directed Medical Research Programs − Melanoma Research Program: Mid-Career Accelerator Award (ME210246; J.E.C.); an award from the National Science Foundation (2217138); a Translational Award Program from the V Foundation (T2023-010); and the Tisch Cancer Institute Cancer Center Support Grant (P30-CA196521). The authors would like to acknowledge the TCI Shared Resources and the Department of Oncological Sciences for research support, and thank Drs. Douglas Green (St. Jude Children’s Research Hospital) and Evripidis Gavathiotis (Albert Einstein College of Medicine) for generously providing *Bid^−/−^Bim^−/−^* and BAX-reconstituted *Bax^−/−^Bak^−/−^* cell lines, respectively.

## AUTHOR CONTRIBUTIONS

Conceptualization: J.D.G. and J.E.C.; methodology: J.D.G., Y.C., A.V.F., and J.N.M.; validation: J.D.G., Y.C., J.N.M., and T.M.S.; investigation: J.D.G., Y.C., M.P.A.L.-V., A.V.F., S.G.B., J.N.M., T.M.S., and N.D.P.; resources: Y.C., M.P.A.L.-V., and M.A.N.; writing: J.D.G. and J.E.C.; visualization: J.D.G.; supervision: Y.S., R.W.K., and J.E.C.; funding acquisition: J.D.G., R.W.K., and J.E.C.

## COMPETING INTERESTS

The authors declare no competing interests.

## METHODS

### Experimental Models

#### Bacterial cell lines

For recombinant protein generation, One Shot^TM^ BL21 (DE3) Chemically Competent *E. coli* were purchased from Invitrogen/Thermo Fisher Scientific (Cat. No. C600003). Cells were grown in BD Difco^TM^ Terrific Broth (Cat. No. BD243820, Thermo Fisher Scientific) media supplemented with 1% glycerol at 37°C with shaking at 220 rpm.

#### Cell lines

*Bax^+/+^Bak^+/+^* and *Bax^−/−^Bak^−/−^* SV40-transformed MEFs were obtained from ATCC (Manassas, VA, USA); *Bim^−/−^Bid^−/−^* SV40-transformed MEFs were provided by Dr. Douglas Green (St. Jude Children’s Research Hospital); *Bax^−/−^Bak^−/−^* double knockout MEFs reconstituted to express wild type or P168G BAX were provided by Dr. Evripidis Gavathiotis (Albert Einstein College of Medicine) and BAX expression was confirmed by western blot and GFP positivity from the pBabe IRES–GFP vector backbone. Cells were cultured in high-glucose DMEM (Cat. No. 10-017-CV, Corning, Manassas, VA, USA) containing 10% FBS (Cat. No. 10438-026, Life Technologies, Carlsbad, CA, USA), 2 mM L-glutamine (Cat. No. 25030-081, Life Technologies, Carlsbad, CA, USA), and 1× Penicillin/Streptomycin (Cat. No. 10378-016, Life Technologies, Carlsbad, CA, USA), and grown in humidified incubators at 37°C with 5% CO2. All cell cultures were maintained in mycoplasm-free conditions as verified by the HEK-Blue Detection Kit (Cat. No. hb-det2, Invivogen, San Diego, CA, USA).

### Recombinant protein and peptides

#### Expression vectors and site-directed mutagenesis

Recombinant human BAX was expressed and purified using an intein-chitin binding domain (CBD) tag from a pTYB1 vector (BAX cDNA inserted into Nde/SapI cloning site, with a C-terminal tag).^49^ BAX mutants (e.g., C62S and C126S double mutant, BAX^2S^; L59C, C62S, C126S, and L162C quadruple mutant, BAX^α8^) were generated by site-directed mutagenesis of the WT construct using oligos provided in the Key Resource Table. Amplification of the *de novo* plasmids was accomplished using CloneAmp HiFi PCR Premix (Cat. No. 639298, Takara Bio, Mountain View, CA, USA) and thermo-cycled as follows: 1× [98°C for 2 minutes]; 30× [98°C for 10 seconds, 55°C for 30 seconds, 72°C for 8 minutes]; and 1× [72°C for 10 minutes]. Parental plasmids were digested by 1 μl DpnI enzyme (Cat. No. R0176, New England Biolabs, Ipswich, MA, USA) at 37°C for 15 minutes, followed by purification using the QIAquick PCR Purification Kit (Cat. No. 28104, Qiagen, Hilden, Germany). Sequences were verified by Genewiz Sanger sequencing using the Universal T7 primer (5’-TAATACGACTCACTATAGGG-3’).

#### Recombinant BAX expression and purification

One Shot™ BL21 (DE3) chemically competent *E. coli* (Cat. No. C601003, Thermo Fisher Scientific) were transformed with the pTYB1-BAX construct in LB broth and grown on agar plates supplemented with 100 μg/mL carbenicillin at 37°C. Starter cultures were grown in Terrific Broth (TB) supplemented with 0.4% glycerol and 100 μg/mL carbenicillin at 30°C for 14–16 hours to an optical density at 600 nm (OD_600_) of approximately 1.5; then diluted with 4× volume of TB broth and expanded at 37°C for 3–4 hours until the culture achieves a target OD_600_ of 2–3. Recombinant BAX expression was induced with 1 mM IPTG at 30°C for 6 hours. Bacteria were pelleted, resuspended in lysis buffer (50 mM K_2_PO_4_, 50 mM NaH_2_PO_4_, 500 mM NaCl, 1 mM EDTA, 0.1 mM AEBSF) supplemented with a Pierce protease inhibitor tablet (Cat. No. A32965, Thermo Fisher Scientific), and lysed with a probe sonicator (e.g., Dismembrator 505, Thermo Fisher Scientific) for 20 minutes. Lysates were centrifuged at 42,000 ×*g* at 4°C for 1 hour to pellet debris and recombinant intein-CBD-tagged BAX was captured from the supernatant by chitin affinity chromatography at 4°C according to the manufacturer’s instructions (Cat. No. S5551, New England Biolabs, Ipswich, MA, USA). On-column intein cleavage of the CBD tag was achieved by incubation with DTT (50 mM) at 4°C for at least 16 hours, followed by elution in Gel Filtration Buffer (20 mM HEPES [pH 7.4], 150 mM NaCl). Recombinant BAX protein (now without a tag or exogenous residues) was further purified by size-exclusion chromatography using Gel Filtration Buffer on a HiLoad 16/600 or 26/600 Superdex 200 pg column (Cat. No. 28989335, Cytiva) using an ÄKTA pure™ 25 L1 (Cat. No. 29018225, Cytiva) at 4°C according to the manufacturer’s instructions. Fractions containing BAX protein were combined and concentrated using Amicon Ultra-4 centrifugal filter (Cat. No. UFC801024, Millipore Sigma) to a stock concentration of ∼30 μM, prior to snap freezing with liquid nitrogen, and storage at –80°C. A detailed version of this protocol has been published.^61^

#### Oxidizing disulfide tether BAX mutants

For the BAX^α8^ mutant, the nascent protein was purified as the reduced form due to incubation with DTT during on-column intein cleavage; to generate a disulfide tether (“locked BAX^α8^”), BAX protein was oxidized in 1 mM Dichloro(1,10-Phenanthroline) Copper (II) (Cat. No. 362204, Millipore Sigma-Aldrich) at 4°C for 15 minutes, purified by Superdex 75 Increase 10/300 GL column (Cat. No. 29148721, Cytiva), and only monomeric species were pooled and concentrated to make protein stocks.

#### Other reagents

Additional recombinant proteins and peptides were purchased from commercial sources: 5-TAMRA labeled BAK-BH3 (Cat. No. AS-64590, AnaSpec); recombinant Bcl–xL^ΔC^, aa 2-212 (Cat. No. 894-BX, R&D Systems); BIM-BH3, Peptide IV (Cat. No. AS-62279, AnaSpec).

### Lipid aldehyde preparation

2-trans-hexadecenal (2t–hexadecenal) was purchased in powder form (Cat. No. 857459P, Avanti Polar Lipids) and reconstituted in pre-warmed (37°C) anhydrous DMSO (Cat. No. D12345, Thermo Fisher Scientific) to a concentration of 50 mM using a Hamilton syringe. The reconstituted 2t–hexadecenal was aliquoted into single-use glass aliquots and stored at -80°C. To avoid precipitation/coagulation of 2t–hexadecenal, stocks were serially diluted first in pre-warmed DMSO, then diluted into pre-warmed aqueous assay buffer to use as a stock for the experiment. This was typically accomplished as follows: 2.5 μl of 50 mM 2t–hexadecenal stocks were two-fold diluted four times in DMSO (to 40 μl of 3.13 mM), then diluted twice in the selected assay buffer (first with 40 μl, then with 45 μl) to achieve a working stock of 125 μl of 1 mM 2t–hexadecenal in 32% DMSO/buffer. Working stocks of 1 mM 2t–hexadecenal were placed in a water bath sonicator for 15 minutes to aid in solubilization and then further diluted to the desired concentration and acceptable DMSO background for experiments. For cell death experiments, 2t–hexadecenal was prepared in 0% FBS media to prevent binding to serum proteins.

Hexadecanal (16:0 aldehyde) was purchased as 1 mg/ml (4.16 mM) in methylene chloride solution (Cat. No. 857458M, Avanti Polar Lipids), aliquoted into 25 μl, and stored at -80°C. Prior to use, hexadecanal was first diluted with pre-warmed DMSO to a concentration of 3 mM, then diluted to 1 mM with pre-warmed assay buffer of choice, sonicated for 15 minutes, and then further diluted for experiments. The remaining alkenals was purchased in liquid form (see Key Resource Table), and their molar concentrations were calculated based on their density, purity, and molecular weight. Short-chain alkenals (2t–5 – 2t–10) were diluted with DMSO to a stock concentration of 1 M, aliquoted, and stored at -80°C. Prior to use, stocks were further diluted with DMSO to 320 mM and then in assay buffer to a final concentration of 1 mM with 32% DMSO (v/v). Long-chain alkenals (2t–11 – 2t–13) were more likely to coagulate and thus were prepared following the exact same procedure as the 2t–hexadecenal preparation.

### Real-time live-cell detection of apoptosis (SPARKL)

#### Cell culture and treatments

MEFs were seeded at 2–4 × 10^3^ cells per well in 96-well tissue culture treated plates and left to adhere for 18–24 hours. Prior to live-cell imaging, growth media was replaced with phenol red-free DMEM as follows: first, 50 μl (50% total volume) of 0% FBS DMEM containing 2t–hexadecenal treatments was added to wells for 15−30 minutes; next, 50 μl of 10% FBS DMEM containing additional treatments and fluorescently-labelled Annexin V (50 nM or ∼1.8 ng/μl) was added to wells. Recombinant Annexin V was generated in-house as previously described.^20,62,63^ Immediately following treatments, plates were subjected to real-time fluorescence microscopy, automated data collection, and analysis.

#### Data acquisition and event detection with an IncuCyte ZOOM

SPARKL experiments performed using an IncuCyte ZOOM (Model 4459, Sartorius, Göttingen, Germany) were housed in a humidified tissue culture incubator maintained at 37°C with 5% CO_2_. Images were collected every 2 hours for 24 hours using a 10× objective, and a single field of view was collected per well. Bright field and green channels (Ex: 440/80 nm; Em: 504/44 nm; acquisition time: 400 ms) were collected at 1392 × 1040 pixels at 1.22 μm/pixel. Automated event detection was accomplished using the ZOOM software (v2018A) and user-defined processing definition using images collected using the relevant cell lines and fluorescent reporters. Processing definition settings were as follows: Parameter, Top-Hat; Radius (μm), 25; Threshold (RFU), 3; Edge Sensitivity, -30; Area (μm^2^), >100. Kinetic data are graphed as the calculated events per well metric provided by the ZOOM software. The y-axis scale was determined for each experiment using parallel internal control treatments to assess maximal apoptotic death. Data are the mean of replicates and are representative of at least three repeated and reproduced assays. A detailed explanation of this method^20^ and protocol^64^ have been published.

#### Data acquisition and event detection with a Cytation 7

SPARKL experiments performed using a Cytation 7 equipped with an inverted imager (Model CYT7UW-SN, BioTek/Agilent, Santa Clara, CA, USA) tethered to a BioSpa 8 automated incubator (Model BIOSPAG-SN, BioTek/Agilent, Santa Clara, CA, USA), where cells were maintained in a humidified environment at 37°C with 5% CO_2_. Bright field and red channel images were collected every 2 hours for at least 24 hours using a 10× objective with a 75% wide field of view crop (1045 μm × 1045 μm), and a single field of view was collected per well using a laser autofocus (Part No. 1225010). Bright field images were acquired with the following settings: LED intensity, 7; Integration Time, 100 msec; Camera Gain, 19. Red channel images (Ex: 586/18 nm; Em: 647/57, Part No. 1225102) were acquired with the following settings: LED intensity, 8; Integration Time, 20 msec; Camera Gain, 19. Automated red event detection was accomplished using the Gen5 software (v3.12) and a cellular analysis data reduction profile with the following settings: Threshold, 5000 RIU; Background, Dark; Split Touching Objects, yes; Fill holes, yes; Background Flattening Size, auto (270 μm); Image Smoothing Strength, 0; Background Percentage, 5%; Minimum Object Size: 5 μm; Maximum Object Size: 90 μm. Kinetic data are graphed as events per image. The y-axis scale was determined for each experiment using parallel internal control treatments to assess maximal apoptotic death. Data are the mean of replicates and are representative of at least three repeated and reproduced assays.

### Large unilamellar vesicle (LUV) permeabilization assays

#### LUV composition and preparation

LUVs and permeabilization assays were prepared as similarly described.^9,65^ Briefly, chicken egg phosphatidylcholine (Cat. No. 840051C, Avanti Polar Lipids), chicken egg phosphatidylethanoloamine (Cat. No. 840021C, Avanti Polar Lipids), porcine brain phosphatidylserine (Cat. No. 840032C, Avanti Polar Lipids), bovine liver phosphatidylinositol (Cat. No. 840042C, Avanti Polar Lipids), and cardiolipin (18:1) (Cat. No. 710335C, Avanti Polar Lipids) were combined at a ratio of 48:28:10:10:4 (5 mg total), dried under N_2_ gas, and resuspended in 500 μl LUV buffer (10 mM HEPES [pH 7], 200 mM KCl, 5 mM MgCl_2_, 0.2 mM EDTA) containing a polyanionic dye (12.5 mM ANTS: 8-aminonaphthalene-1,3,6-trisulfonic acid) and cationic quencher (45 mM DPX: p-xylene-bis-pyridinium bromide) using a water bath sonicator. Unilamellar vesicles were formed by 31 extrusions of the suspension through 1.0 μm polycarbonate membranes (Cat. No. 610010, Avanti Polar Lipids, Alabaster, AL, USA). The unincorporated DPX and ANTS were removed by using a 10 ml Sepharose S-500 gravity flow column. LUVs were used within 2 weeks of preparation to avoid significant liposome degradation.

#### LUV assays

Working stocks of recombinant proteins, peptides, and lipids were prepared in LUV buffer at 4× their intended final concentrations. LUVs from the preparation stock (20×) were diluted five-fold in LUV buffer to generate a working stock (4×) for assays (equivalent to each sample receiving 5 μl undiluted LUVs). In a black opaque 96-well plate, lipids or peptide titrations were generated via in-well serial dilutions and buffer was added to any control wells requiring volume compensation. Samples were assayed in 100 μl total volume with 25 μl of each component sequentially added to achieve desired 1× concentrations. A typical assay was prepared as follows: 25 μl of titrant is prepared in the well; then 25 μl of peptide, reagent, or buffer is added; then 25 μl of recombinant BAX; and finally, 25 μl of diluted LUVs. Plates were immediately subjected to fluorescence analyses for 90 minutes using either a Synergy H1 or Cytation 5 multi-mode microplate reader (BioTek/Agilent, Santa Clara, CA, USA) using parameters listed in the table below. Kinetic data represents the mean of triplicate samples and are representative of at least three repeated and reproduced assays using separate protein aliquots and LUV preparations. Every assay included a 1% CHAPS positive control to determine maximum signal. Normalized endpoint data (% permeabilization) were calculated in Prism (Graphpad) using the minimum value of the buffer control (as 0%) and the mean value of LUVs solubilized in 1% CHAPS (as 100%) measured during the entire assay (see Equation 1). Appropriate concentrations of BAX were determined by extensive protein titrations for each preparation of recombinant protein and LUVs; typically the highest BAX concentration exhibiting minimal permeabilization was selected for subsequent assays to ensure adequate signal and dynamic range for activation studies.

<u>Equation 1: LUV data normalization</u>

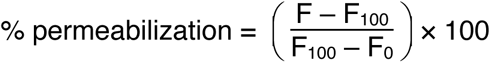

### Microscale thermophoresis

MST experiments were carried out on a NT.115 (NanoTemper GmbH, Munich, Germany) instrument and performed in MST buffer (50 mM Tris-HCl pH 7.4, 150 mM NaCl, 10 mM MgCl_2_, 0.05% Tween-20). K_D_ values were calculated using the NanoTemper software. The recombinant human BAX protein was labeled with Alexa Fluor-647 NHS ester by incubating 60 μM of the fluorescent dye with a 20 μM of BAX protein for 30 minutes in the dark. The labeling mixture was subsequently applied to a G-25 gravity flow column (GE Healthcare) that had been equilibrated with MST buffer. After elution from the G-25 column, the concentration of the labeled protein was quantified spectrophotometrically, snap frozen in liquid nitrogen, and stored as single use aliquots at -80°C. Labeled BAX protein was treated with 0.002% CHAPS to prevent activation-induced BAX aggregation upon ligand binding. Curve fitting was conducted using Prism (Graphpad) and only applied to X values from 5−35 seconds using Equation 2, and the K constant was calculated using Equation 3.

<u>Equation 2: One-phase decay function for MST data fitting</u>

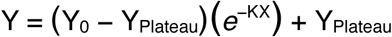

<u>Equation 3: Decay constant function</u>

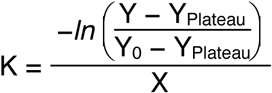

### Liquid chromatography-mass spectrometry (LC-MS)

#### Sample preparation and handling

Working stock of recombinant BAX protein and 2t–hexadecenal were prepared in Gel Filtration Buffer. 30 μl of 5 μM recombinant BAX (∼3 μg) was treated as indicated and incubated at 37°C for 2 hours. Samples were then reduced in 8 M urea buffer (with 50 mM NH_4_HCO_3_, 10 mM TCEP) at 57°C for 1 hour, and alkylated in the dark with 30 mM iodoacetamide for 30 minutes at 25°C. Samples were then digested using 1:100 (w/w) trypsin at 37°C for 4 hours, with an additional round of 1:100 trypsin at 37°C for 16–18 hours. After proteolysis, the peptide mixtures were desalted by self-packed stage-tips or Sep-Pak C18 columns (Waters) and analyzed with a Vanquish Neo UHPLC System that is coupled online with a Orbitrap Exploris 480 mass spectrometer (Thermo Fisher). Briefly, desalted Nb peptides were loaded onto an analytical column (C18, 1.7 μm particle size, 150 μm × 5 cm; IonOpticks) and eluted using a 20 minute liquid chromatography gradient (3–10% B, 0–2 minutes; 10–40% B, 2–12 minutes; 40–80% B, 12–14 minutes; 3% B, 14–20 minutes). Mobile phase A consisted of 0.1% formic acid (FA), and mobile phase B consisted of 0.1% FA in 80% acetonitrile (ACN). The flow rate was 1 μl/minute. The Orbitrap Exploris 480 instrument was operated in the data-dependent mode, where the top 20 most abundant ions (mass range 200–1800, charge state 2–6) were fragmented by high-energy collisional dissociation (HCD). The target resolution was 120,000 for MS and 15,000 for tandem MS (MS/MS) analyses. The maximum injection time for MS/MS was set at 100 ms.

#### Proteomic data analysis

Raw data collected by LC-MS was searched against the Uniprot reviewed Human protein sequences database retrieved on 01 March 2024 with decoys and common contaminants appended using FragPipe (v22.0). A labile search was performed in MSFragger^66^ without diagnostic and Y ions specified. Mass offsets were set restricted to cysteines. The offset list was set at 0 (no modification) and 238.22967 (monoisotopic mass of 2t–hexadecenal) and replacing the fixed cysteine carbamidomethylation with a variable one. “Write calibrated MGF” was turned on for the PTM-Shepherd^67^ diagnostic feature mining module, and “Diagnostic Feature Discovery” in PTM-Shepherd was enabled with default parameters. Peptide and protein levels were performed using label-free quantification (LFQ) algorithms in IonQuant^68^. After MS search with MSFragger, raw files and identification files were imported into PDV^69^ (https://github.com/wenbostar/PDV) for MS spectra annotation.

### Thermal Shift Differential Scanning Fluorimetry

Working stock of recombinant BAX proteins, SYPRO Orange dye (Cat. No. S6650, Thermo Fisher Scientific), and 2t–hexadecenal were prepared in gel filtration buffer. The assay was performed in 100 μl 96-well PCR plates using a total assay volume of 50 μl. 20 μl of 2.5× 2t–hexadecenal titrations were generated via in-well serial dilutions and buffer was added to any control wells requiring volume compensation. 20 μl of 2.5 μM BAX and 10 μl of 20× SYPRO Orange were sequentially added to the wells, thoroughly mixed by pipetting, and the plate was centrifuged to recollect sample in the bottom of the well and remove any trapped bubbles. The PCR plate was then sealed with optically clear adhesive sheet and subjected to fluorescence spectroscopy using an Applied Biosystems ViiA7 real-time PCR instrument (Thermo Fisher Scientific). Temperature started at 25°C and increased to 95°C at a rate of 1% per minute. Data was collected as normalized fluorescence at each step of the thermal ramp and the melting temperature was determined as the maximum first derivative value. Data shown are the average of at least four replicates and error bars denote the SEM.

### Fluorescence polarization ligand assay to monitor BAX early-activation (FLAMBE)

#### Assay setup

Working stocks of recombinant BAX protein, peptides, and lipids were prepared in 0.5× PBS buffer at 4× their intended final concentrations. 5-TAMRA labeled BAK-BH3 peptide (Cat. No. AS-64590, AnaSpec) was diluted to a 200 nM working stock (4×). In a black opaque 96-well plate, lipids or peptide titrations were generated via in-well serial dilutions and buffer was added to any control wells requiring volume compensation. Samples were assayed in 100 μl total volume with 25 μl of each component sequentially added to achieve desired 1× concentrations. A typical assay was prepared as follows: 25 μl of titrant is prepared in the well; then 25 μl of peptide, reagent, or buffer is added; then 25 μl of recombinant BAX; and finally, 25 μl of BAK^TAMRA^. Plates were immediately subjected to spectrometry analyses for 60 minutes using a Synergy H1 multi-mode microplate reader equipped with a polarization filter (BioTek/Agilent, Santa Clara, CA, USA) using parameters listed in the table below. Polarization (expressed as milli-Polarization (mP) units) is derived from measured parallel and perpendicular emission intensities (I) and was calculated by the Gen5 software (BioTek/Agilent); Polarization can also be calculated manually using Equation 2. Kinetic data represents the mean of triplicate samples and are representative of at least three repeated and reproduced assays using separate protein aliquots. Appropriate concentrations of BAX and peptides were determined by extensive protein titrations for each preparation of recombinant protein; typically, the highest BAX concentration exhibiting no BAK^TAMRA^ dissociation was selected for subsequent assays to ensure adequate signal and dynamic range for activation studies. Similarly, concentrations of direct activators were selected from extensive titration experiments to determine the appropriate concentration for each experimental setup.

#### Kinetic FP data parameterization

Parameterized FLAMBE data was derived from the average of replicates to reduce data noise. For each condition, Tmax was identified as the timepoint with the highest Polarization and EP was the endpoint Polarization value recorded. Tmax of the BAK^TAMRA^ control or any sample exhibiting no binding kinetics during the assay was set to 0 to avoid misidentification due to noise. Each parameterized metric was normalized to metrics from the BAK^TAMRA^ and BAX control conditions, as 0 and 1 respectively. The shift magnitude of parameterized FLAMBE data was calculated using the distance between conditions without and with BIM-BH3 treatment (see Equation 3). Area under the curve (AUC) was calculated using Prism (Graphpad) and normalized to the controls. A detailed explanation of this method^8^ and protocol^70^ have been published.

#### Competition FP assay

BCL–xL competitive binding FP assays were conducted similarly to FLAMBE assays with some modifications. BCL–xL, 2t–hexadecenal, and BAK^TAMRA^ were prepared as 4× working stocks in a modified FP assay buffer (20 mM sodium phosphate buffer [pH 7.4], 50 mM NaCl, 1 mM EDTA, 0.05 % pluronic F-68).^71^ 2t–hexadecenal titrations were prepared in-well and diluted with 25 μl buffer; separately, recombinant BCL–xL and BAK^TAMRA^ stocks are combined at a 1:1 volume ratio (resulting in a 2× stock) and 50 μl is added to sample wells immediately followed by spectrometry.

<u>Equation 2: Polarization calculation</u>

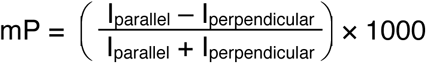

<u>Equation 3: Distance equation for parameterized FLAMBE data</u>

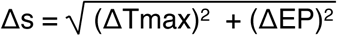

### Fluorescent size-exclusion chromatography

Alexa Fluor™ 488 NHS ester (AF488)-labeled BAX was generated according to the manufacturer’s instructions (Thermo Fisher Scientific, Cat. No. A20000) at a 20:1 dye:protein molar excess. Excess dye was removed by gel filtration with a Superdex 75 Increase 10/300 GL column (Cat. No. 29148721, Cytiva) equilibrated with Gel Filtration Buffer using an ÄKTA pure™ 25 L1 (Cat. No. 29018225, Cytiva) at 4°C according to the manufacturer’s instructions. 0.5 ml fractions were collected and fractions containing monomeric AF488-labelled BAX were combined, quantified, aliquoted, and stored at -80°C. For experiments, 600 μl of 400 nM 488-labelled BAX was treated as indicated at 25°C for 1 hour and protected from light. Following treatment, BAX was subjected to size exclusion chromatography using the above parameters. For detection, 0.2 ml of eluate from the indicated fractions was transferred to a black 96 well-plate and analyzed for fluorescence by a Cytation 5 multi-mode plate reader (BioTek/Agilent, Santa Clara, CA, USA) using the following parameters: Excitation, 495/10 nm; Emission, 520/10 nm; Gain (volage), 100.

### 2D#HSQC NMR spectroscopy

^15^N-labeled BAX protein was prepared at 40 μM in 15 mM sodium phosphate (pH 7.0), 50 mM NaCl. Data were acquired on Bruker 600-MHz spectrometer equipped with a cryoprobe. Two-dimensional ^1^H-^15^N correlation spectra were acquired at 25°C using standard Bruker pulse sequences using 128 scans, 2,048 × 200 complex points and spectral windows of 14 p.p.m. × 35 p.p.m. in the ^1^H and ^15^N dimensions, respectively. Spectra were processed using TOPSpin (Bruker Biospin, MA, USA) and analyzed with CARA software^72^ (cara.nmr.ch). The weighted average chemical shift perturbation (CSP) difference (Δδ) was calculated using Equation 1. The absence of a value indicates no CSP difference, the presence of a proline, or a residue that is overlapped or missing and therefore not used in the analysis. The significance threshold for CSPs was calculated based on the average chemical shift across all measurable residues plus 1 or 2 standard deviations, in accordance with standard methods. Mapping of CSP data onto the BAX structure (PDB: 1F16) was performed with PyMOL (Schrödinger, LLC).

<u>Equation 4: CSP difference equation</u>

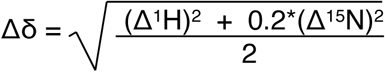

### *In silico* docking and molecular modeling

#### Unbiased docking simulations

The molecular interactions between the alkenals and BAX were modeled using the EADock DSS version of the SwissDock webserver (www.swissdock.ch).^73,74^ The structure PDB files and the alkenal mol2 files were obtained from the Protein Data Bank and the ZINC database, respectively (see Table 3). Output files were split into individual clusters and visualized using PyMOL (Schrödinger, LLC). Quantification of alkenal binding was accomplished by manual inspection of each cluster group and the total number of binding modality outputs for each job. Cavity determination and visualization was performed with PyMOL using a cavity radius of 3 and a cavity cutoff of -5.5; additional spaces were excluded to enhance visualization.

**Table 1:** Parameters for LUV permeabilization assays

|  |  |
| --- | --- |
| Reader | BioTek Synergy H1 or Cytation 5 multi-mode plate reader |
| Read interval | 55 seconds |
| Total read time | 90 minutes |
| Protocol | 37°C; 2 second linear shake; read |
| Excitation | 355/20 nm |
| Emission | 520/20 nm |
| Gain (voltage) | 100 |
| Optics position | Top |
| Read height | 5.5 mm |
| Assay plate | black polystyrene 96-well (Corning #3915) |
| LUV buffer | 10 mM HEPES (pH 7), 200 mM KCl, 5 mM MgCl<sub>2</sub>, 0.2 mM EDTA |

**Table 2:** Parameters for FP assays

|  |  |
| --- | --- |
| Reader | BioTek Synergy H1 multi-mode plate reader |
| Filter Cube | red FP polarizer filter set (#8040562) |
| Read interval | 30 seconds |
| Total read time | 60 minutes |
| Protocol | 25°C; 2 second double orbital shake; read |
| Excitation | 530/25 nm |
| Emission | 590/35 nm |
| Mirror | 570 nm |
| Gain (voltage) | 50 |
| Optics position | Top |
| Read height | 7.0 mm |
| Assay plate | black polystyrene 96-well (Corning #3915) |
| FLAMBE buffer | 0.5× PBS |
| Anti-apoptotic FP buffer | 20 mM phosphate buffer (pH 7.4), 50 mM NaCl, 1 mM EDTA, 0.05 % pluronic F-68 |

**Table 3:**
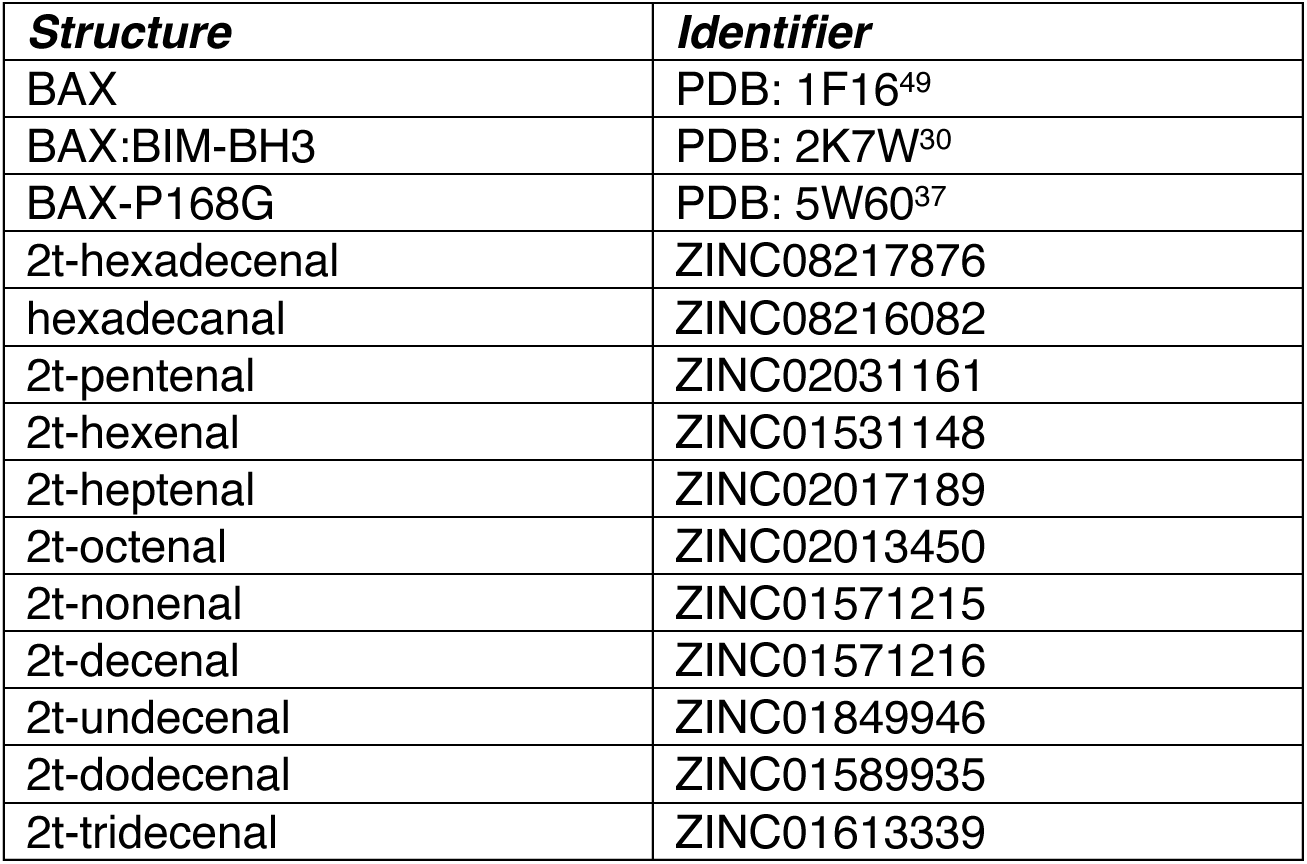
List of resources for structural investigations

| **Structure** | **Identifier** |
| --- | --- |
| BAX | PDB: 1F16<sup>49</sup> |
| BAX:BIM-BH3 | PDB: 2K7W<sup>30</sup> |
| BAX-P168G | PDB: 5W60<sup>37</sup> |
| 2t-hexadecenal | ZINC08217876 |
| hexadecanal | ZINC08216082 |
| 2t-pentenal | ZINC02031161 |
| 2t-hexenal | ZINC01531148 |
| 2t-heptenal | ZINC02017189 |
| 2t-octenal | ZINC02013450 |
| 2t-nonenal | ZINC01571215 |
| 2t-decenal | ZINC01571216 |
| 2t-undecenal | ZINC01849946 |
| 2t-dodecenal | ZINC01589935 |
| 2t-tridecenal | ZINC01613339 |

#### Virtual mutagenesis and molecular docking

Virtual mutagenesis of BAX structures was performed using the mutagenesis wizard in PyMOL (v3.1, Schrödinger, LLC) and selecting rotamers that did not conflict with surrounding sidechains; the resulting structures were utilized in downstream investigations. NMR-guided molecular docking of 2t–hexadecenal on BAX was performed using Glide (v2023–1 build 128, Schrödinger, LLC) with extra precision (XP) and a receptor grid generated in the center of the BAF (13 × 13 × 13 Å inner box, 10 Å outer box), with no additional constraints. Wild type and mutant structures had α8 residues (157−163) removed and were prepared using the Protein Prep Wizard, assigning partial charges with EPIK (Schrödinger, LLC), and aligned against the 1F16 structure to ensure consistency of BAF coordinates for grid generation. The 2t–hexadecenal structure was converted to 3D and prepared for docking using LIGPREP (v2023–1 build 128, Schrödinger, LLC). The lowest energy docking pose is visualized, was representative of all output poses, and various scoring metrics are reported. Interaction fingerprints were generated for residues exhibiting overall interaction with any of the output poses. WT and P168G BAX interaction comparisons were calculated as the fraction of total poses interacting with a residue, relative to the WT:2t–hexadecenal interaction, and color-coded for visualization in PyMOL (Schrödinger, LLC). Interaction diagrams were generated and exported from MAESTRO (Schrödinger, LLC) and then redrawn in Inkscape (www.inkscape.org) to aid in visualization.

### SDS-PAGE and Western Blotting

*Bax^−/−^Bak^−/−^* MEFs, or MEFs reconstituted to express WT or P168G BAX, were harvested with 0.25% Trypsin-EDTA and pelleted at 800× g for 5 minutes. Cells were lysed with 1× RIPA lysis buffer supplemented with protease inhibitors (HALT Tablet, Cat. No. 87786) and phosphatase inhibitors (ApexBio, Cat. No. k1015b) on ice for 20 minutes, pelleted at 21,000× g for 10 minutes at 4°C, and supernatant was collected for protein quantification by BCA assay (Cat. No. 23225, Thermo Fisher Scientific, Waltham, MA, USA) according to the manufacturer’s protocol. Sample concentrations were equilibrated in lysis buffer, combined with 4× Laemmli sample loading buffer supplemented with DTT, subjected to SDS-PAGE in a 12.5% polyacrylamide gel, and transferred to nitrocellulose by standard western blot conditions. Membranes were blocked in 5% milk in 1× TBS buffer supplemented with 0.1% Tween-20, incubated with primary antibodies (1:500-1000 in blocking buffer; incubated for 16−18 hours at 4°C) and secondary antibodies (1:4000 in blocking buffer; incubated for 1 hour at 25°C), followed by standard enhanced chemiluminescence detection (Cat. No. WBLUF0100, Sigma-Aldrich, St. Louis, MO, USA). Antibodies: BAX, 1:500 dilution (Clone 2D2, Cat. No. sc-20067, Santa Cruz Biotechnology, Dallas, TX, USA); GAPDH, 1:1000 dilution (Clone 1E6D9, Cat. No. 60004, Proteintech, Rosemont, IL, USA); m-IgGk BP-HRP secondary antibody (Cat. No. sc-516102, Santa Cruz Biotechnology, Dallas, TX, USA).

## DATA AND RESOURCE AVAILABILITY

The data supporting the findings of this study are included in this published article (and its supplementary information files). Materials generated in this study are available from the corresponding author upon reasonable request. Structures corresponding to PDB 1F16, 2K7W, and GW60 were used in this study.

## SUPPLEMENTARY INFORMATION

### Supplementary figure titles

**Supplementary Figure 1:**
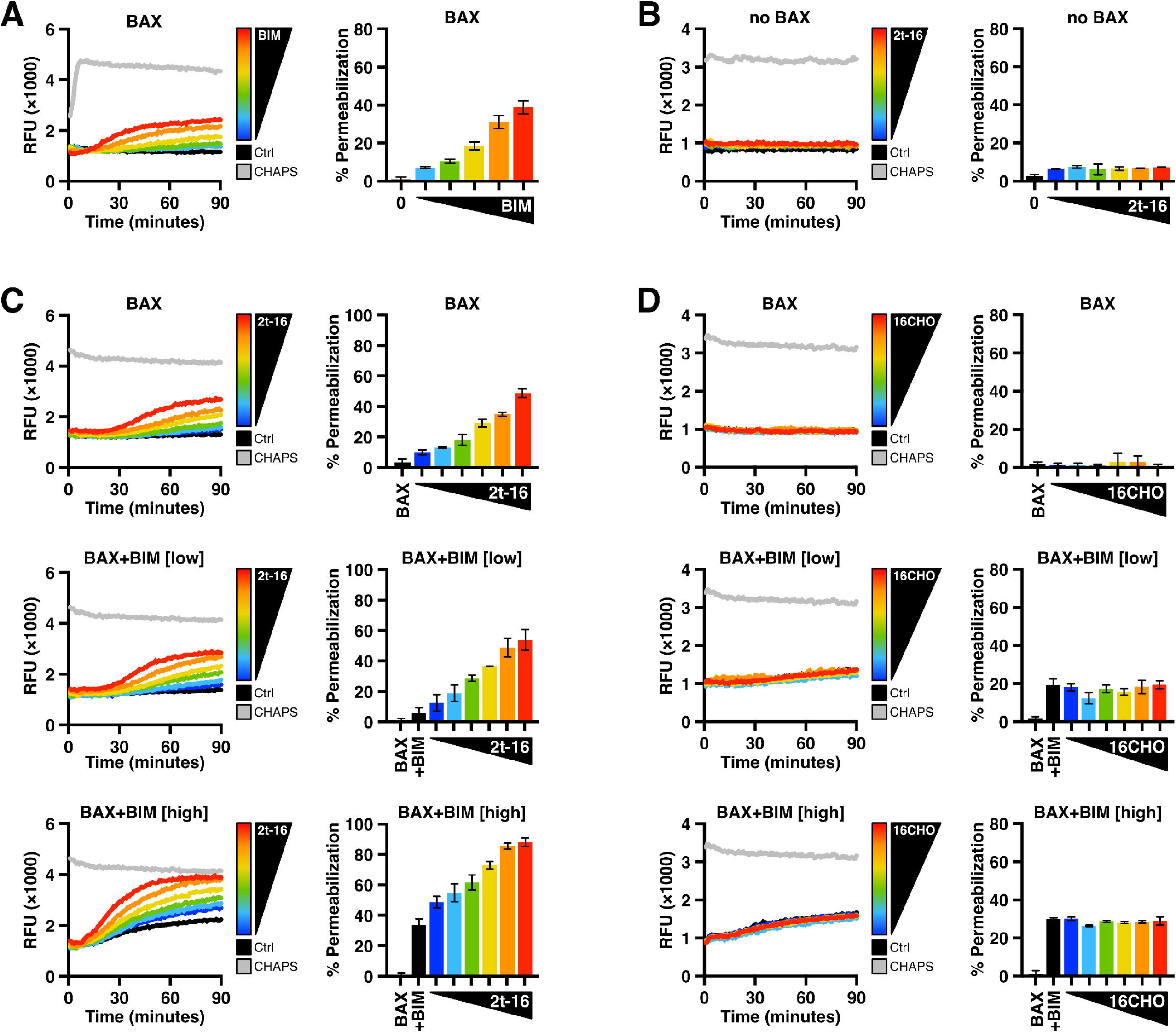
2t–hexadecenal promotes BAX functionalization (Related to Figure 1) **(A–D)** LUV permeabilization studies with recombinant BAX protein treated as indicated and measured at regular intervals for changes in fluorescence as fluorophores are released from compromised liposomes. Left panels: kinetic fluorescence data; right panels: endpoint data normalized to LUV fluorescence and maximal signal generated by LUVs solubilized with CHAPS detergent (grey data). Data are shown as the mean of technical replicates and error bars report SEM. **(A)** BAX protein (100 nM) was activated by BIM–BH3 peptide (0.13–2 μM) and added to LUVs. **(B)** LUVs treated with 2t–16 (16.5–50 μM) in the absence of BAX to confirm no membrane destabilization by 2t–16. **(C)** Data summarized by Figure 1F. LUVs permeabilized by BAX (120 nM) treated with 2t–16 (6.5– 50 μM) ± BIM–BH3 peptide (0.5, 2.5 μM). **(D)** Data summarized by Figure 1I. LUVs permeabilized by BAX (160 nM) treated with 16CHO (6.5–50 μM) ± BIM–BH3 peptide (0.5, 2.5 μM).

**Supplementary Figure 2:**
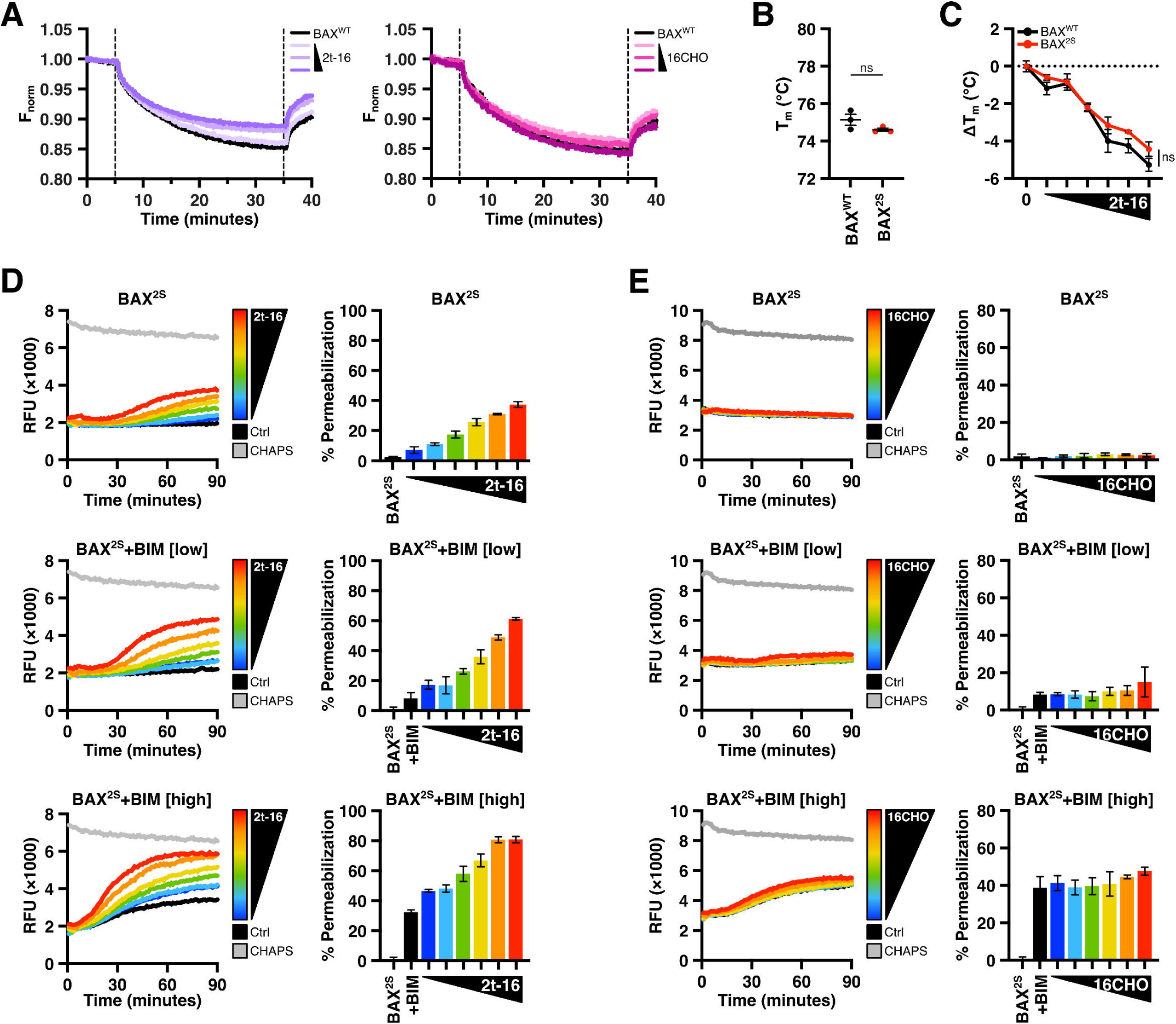
BAX activation by 2t–hexadecenal does not require cysteine residues (Related to Figure 2) **(A)** Alexa Fluor 647-labeled recombinant BAX^WT^ (1 nM) was incubated with CHAPS (0.002%) to inhibit oligomerization, treated as indicated, and subjected to MST. Timetrace thermal shift curves of BAX^WT^ titrated with 2t–16 or 16CHO (0.04, 1.25, 40 μM) report the mean of replicate data. **(B−C)** The melting temperature of BAX^WT^ and BAX^2S^ ± 2t–16 (6.5−50 μM) was measured by thermal shift assay using SYPRO orange and compared. Statistical significance was determined by two-way ANOVA; ns, not significant (*P* > 0.05). **(D–E)** LUV permeabilization studies with recombinant BAX^2S^ protein treated as indicated and measured at regular intervals for changes in fluorescence as fluorophores are released from compromised liposomes. Left panels: kinetic fluorescence data; right panels: endpoint data normalized to LUV fluorescence and maximal signal generated by LUVs solubilized with CHAPS detergent (grey data). Data are shown as the mean of technical replicates and error bars report SEM. **(D)** Data summarized by Figure 2F. LUVs permeabilized by BAX^2S^ (100 nM) treated with 2t–16 (6.5–50 μM) ± BIM–BH3 peptide (0.5, 2.5 μM). **(E)** Data summarized by Figure 2I. LUVs permeabilized by BAX^2S^ (100 nM) treated with 16CHO (6.5–50 μM) ± BIM–BH3 peptide (0.5, 2.5 μM).

**Supplementary Figure 3:**
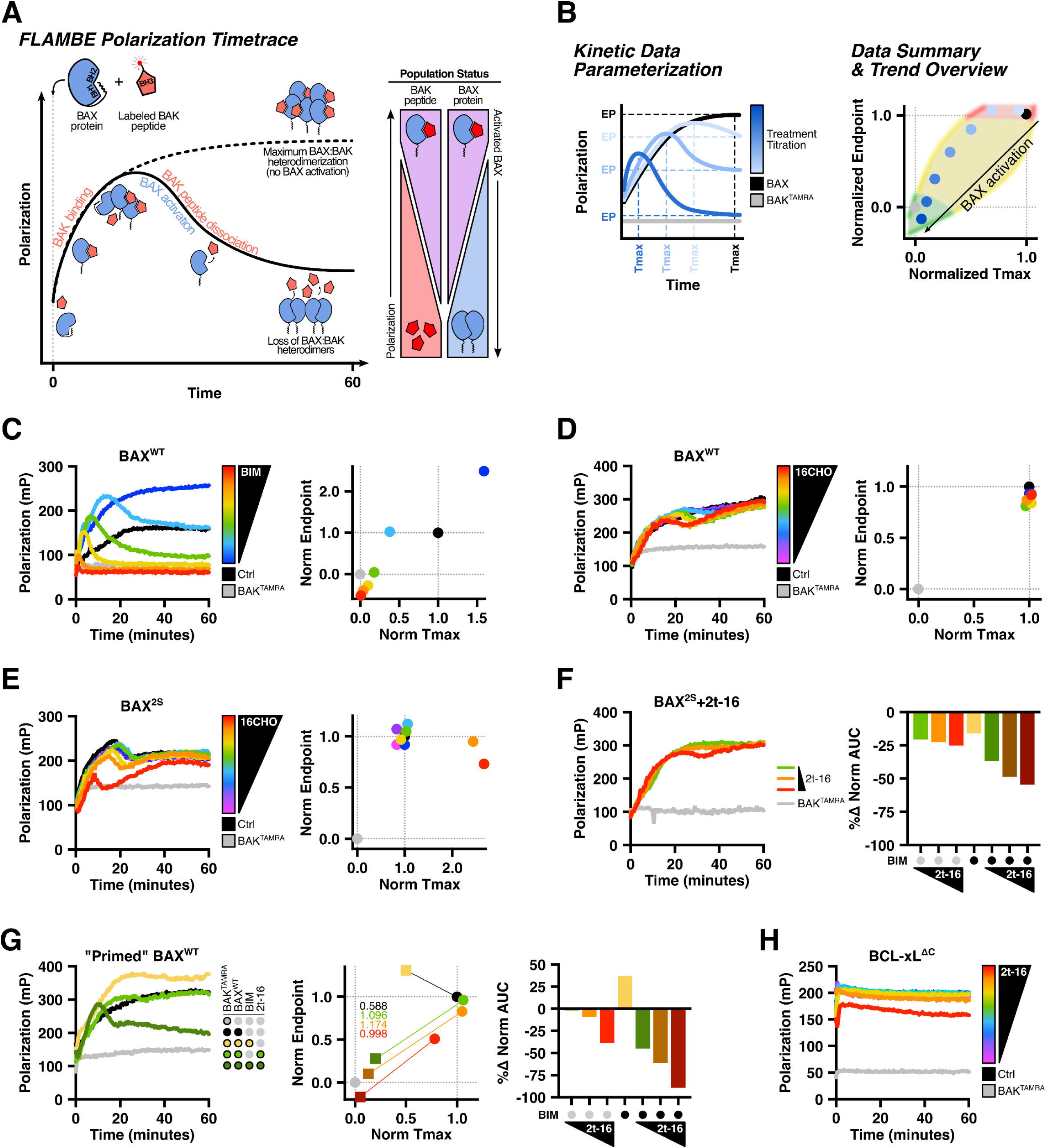
An assay for detecting BAX early-activation steps reveals that 2t–hexadecenal cooperates with BIM-mediated triggering (Related to Figure 3) **(A)** Illustration of BAX and BAK–BH3 interactions within FLAMBE. The reporter is a fluorescently-labeled BAK–BH3 peptide that exhibits changes in Polarization measurements as it is bound by BAX. Over time, the BAX population binds BAK–BH3 peptides resulting in increased Polarization, which eventually plateaus if the entire population forms heterodimers (dotted line). In conditions that activate BAX, activation-induced intramolecular rearrangements within BAX result in the dissociation of the BAK–BH3 peptide and a concomitant decrease in Polarization over time as the population of unbound BAK–BH3 peptide increases (solid line). **(B)** Illustration of FLAMBE data parameterization and analysis. Left: Kinetic Polarization data is collected for a treatment causing BAX activation and exhibiting accelerated kinetics of BAK^TAMRA^ dissociation (blue lines, depicted as a titration exhibiting dose-dependent BAX activation). Kinetic data is parameterized by extracting endpoint Polarization (EP) and time-to-maximum peak (Tmax) for each condition. Right: Parameterized data are normalized to BAK^TAMRA^ and BAX controls (grey and black, respectively) and titrations or separate conditions can be visualized as a two-dimensional plot. Generally, conditions exhibiting minimal or robust activation of the BAX population cluster in the upper-right or lower-left regions, respectively. Conditions plotted above BAX (i.e., EP > 1) form stable non-activating complexes with the BAX:BAK^TAMRA^ heterodimer (red region). **(C)** BAX^WT^ (60 nM) was treated with BIM–BH3 peptide (0.25–2 μM) and subjected to FLAMBE to visualize dose-dependent activation-induced dissociation of BAK^TAMRA^. BIM–BH3 at a low concentration (0.25 μM, dark blue data) demonstrated a stable, non-activating interaction with the BAX:BAK^TAMRA^ complex and exhibited increased Polarization. **(D)** BAX^WT^ (60 nM) was treated with 16CHO (2–50 μM), combined with BAK^TAMRA^ (50 nM), and subjected to FLAMBE. **(E)** Same as in **D** with BAX^2S^ (60 nM). **(F)** Left: BAX^2S^ (60 nM) was treated with three non-activating concentrations of 2t–16 (green: 4.5 μM; orange: 6.5 μM; red: 10 μM). Parameterization of this data is included in Figure 3D. Right: AUC calculated for each condition was normalized to the BAX and BAK^TAMRA^ controls and reported as a percent change from the vehicle-treated BAX condition. **(G)** Left: BAX^WT^ (60 nM) was combined with a non-activating concentrations of BIM–BH3 peptide (0.15 μM) and 2t–16 (4.5 μM), followed by BAK^TAMRA^ (50 nM), and subjected to FLAMBE. Middle: Parameterized FLAMBE data including three concentrations of 2t–16 (green: 4.5 μM; orange: 6.5 μM; red: 10 μM) in the absence or presence of BIM–BH3 (circle and square datapoints, respectively). Annotations report the magnitude of shift between data with and without BIM–BH3. Right: AUC calculated for each condition was normalized to the BAX and BAK^TAMRA^ controls and reported as a percent change from the vehicle-treated BAX condition. **(H)** Fluorescence polarization competition assay with recombinant BCL-xL^ΔC^ protein treated with 2t–16 (2–50 μM) and combined with BAK^TAMRA^.

**Supplementary Figure 4:**
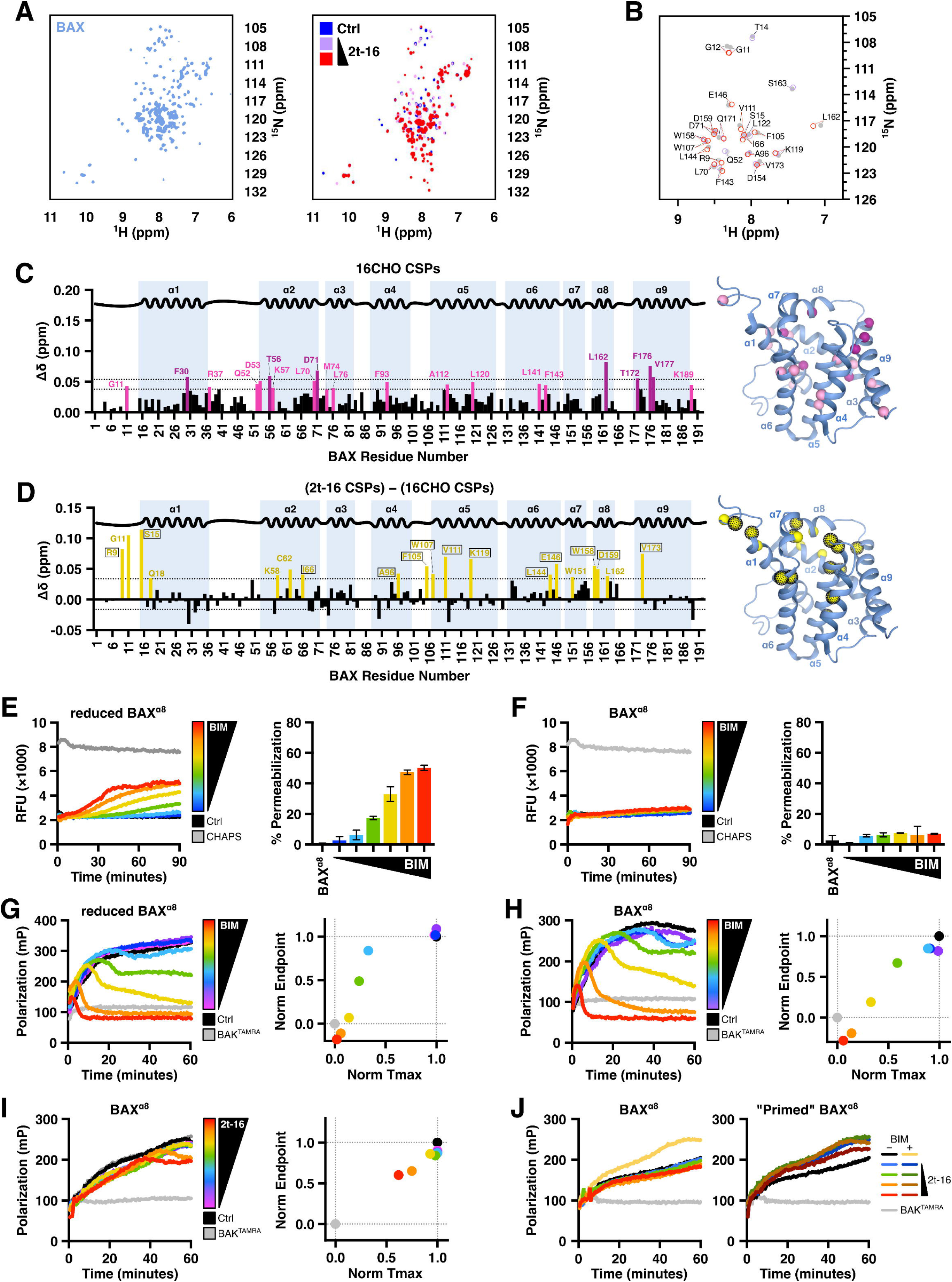
Structural, biophysical, and functional approaches identify α8 mobilization as necessary for 2t–hexadecenal function (Related to Figure 4) **(A)** ^1^H-^15^N HSQC NMR spectra of ^15^N-labeled BAX^WT^ (40 μM) and BAX^WT^ treated with vehicle or 2t–16 (50, 150 μM). **(B)** Plot of BAX residues exhibiting a significant shift in response to incubation with 2t–16. **(C)** Chemical shift perturbations (CSPs) observed in ^15^N-labeled BAX^WT^ incubated with 16CHO (150 μM). Residues exhibiting a shift greater than 1 or 2 standard deviations above the average (dotted lines) are highlighted and indicated on the BAX structure (PDB: 1F16). The absence of a bar indicates no chemical shift difference, the presence of a proline, or the residue that could not be definitively assigned. **(D)** Chemical shift perturbations (CSPs) observed in ^15^N-labeled BAX incubated with 2t–16 (Figure 4A) were subtracted by CSPs observed with 16CHO (Figure **S4C**) and plotted as a function of BAX residues. Residues exhibiting a shift greater than the 1 standard deviation above the average (dotted line) are colored yellow and indicated on the BAX structure (PDB: 1F16). Residues uniquely exhibiting significance in the 2t–16 treatment and not 16CHO are highlighted with a black outline. The absence of a bar indicates no chemical shift difference, the presence of a proline, or the residue that could not be definitively assigned. **(E)** LUVs permeabilized by reduced BAX^α8^ (220 nM) treated with BIM–BH3 peptide (0.2–6.5 μM). Data are shown as the mean of technical replicates and error bars report SEM. **(F)** Same as in **E** with oxidized BAX^α8^ (220 nM) to induce a disulfide tether immobilizing α8. **(G–J)** BAX^α8^ was treated as indicated, combined with a TAMRA-labeled BAK–BH3 peptide (BAK^TAMRA^), and immediately subjected to FLAMBE analysis. Left panels: kinetic Polarization data; right panels: two-dimensional plot of parameterized FLAMBE data comparing Tmax and endpoint Polarization metrics normalized to BAK^TAMRA^ (grey data) and the vehicle-treated BAX control (black data). **(G)** Reduced BAX^α8^ (75 nM) was treated with BIM–BH3 (0.06–1 μM), combined with BAK^TAMRA^ (50 nM), and subjected to FLAMBE. **(H)** Same as in **G** with oxidized BAX^α8^ (250 nM) to immobilize α8. **(I)** Oxidized BAX^α8^ (250 nM) was treated with 2t–16 (3–50 μM), combined with BAK^TAMRA^ (50 nM), and subjected to FLAMBE. **(J)** BAX^α8^ (225 nM) was treated with four non-activating concentrations of 2t–16 (blue: 3 μM; green: 4.5 μM; orange: 6.5 μM; red: 10 μM) in the absence or presence of BIM–BH3, followed by BAK^TAMRA^ (50 nM), and subjected to FLAMBE. Parameterization of this data is included in Figure 4I.

**Supplementary Figure 5:**
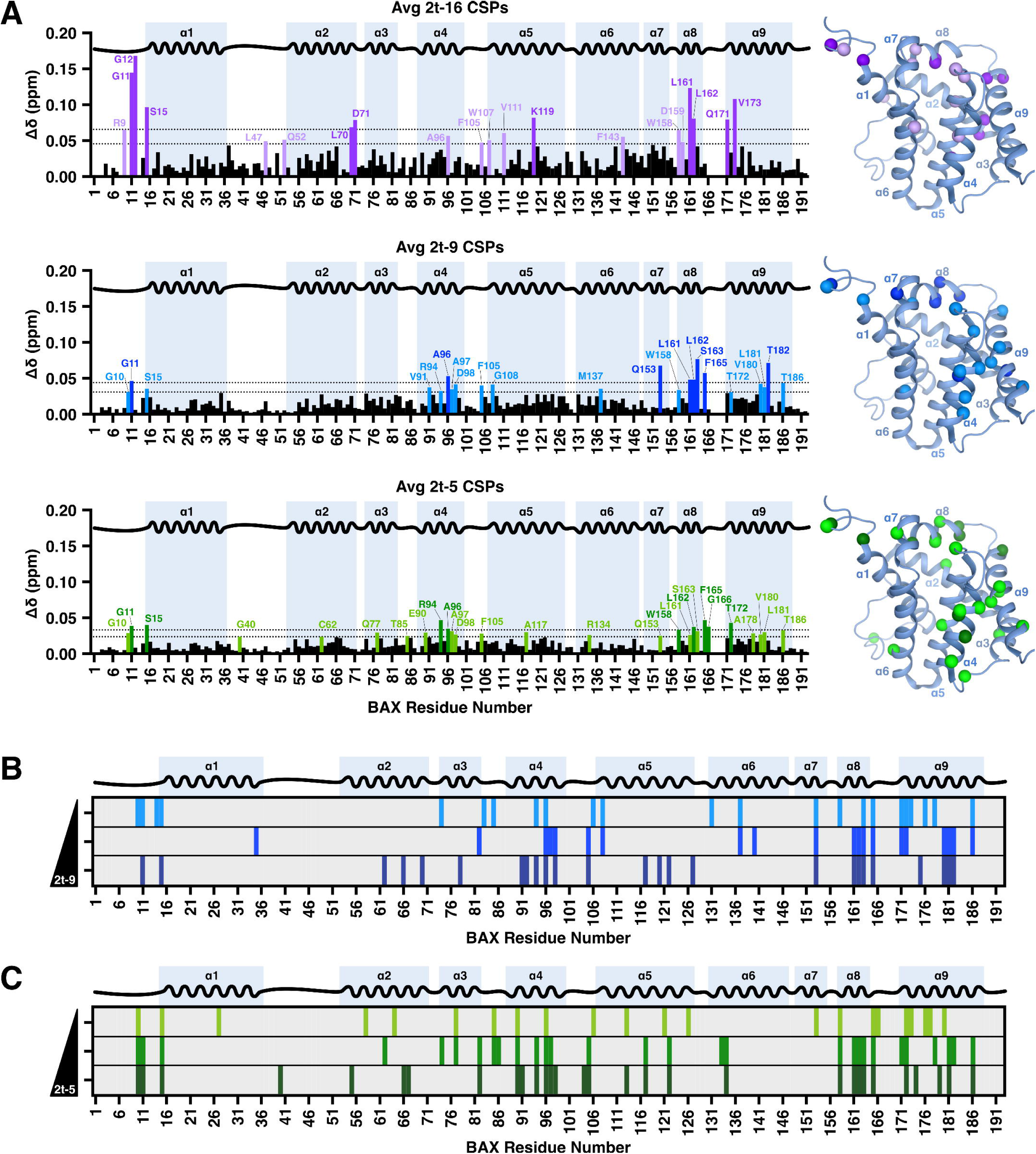
^1^H-^15^N HSQC perturbations indicate that short-chain 2t–alkenals exhibit CSPs outside of the BAF region (Related to Figure 5) **(A)** Chemical shift perturbations (CSPs) observed in ^15^N-labeled BAX incubated with either 2t–16 (50, 150 μM), 2t–9 (0.3, 0.9, 2.7 mM), or 2t–5 (0.3, 0.9, 2.7 mM) averaged across concentrations. Residues exhibiting a shift greater than 1 or 2 standard deviations above the average (dotted lines) are indicated in light and dark colors, respectively, and indicated on the BAX structure (PDB: 1F16). The absence of a bar indicates no chemical shift difference, the presence of a proline, or the residue that could not be definitively assigned. **(C–D)** Residues exhibiting significant CSPs for each concentration (0.3, 0.9, 2.7 mM) of 2t–9 or 2t–5. Highlighted residues exhibited shifts greater than 1 standard deviation above the average of measurable shifts.

**Supplementary Figure 6:**
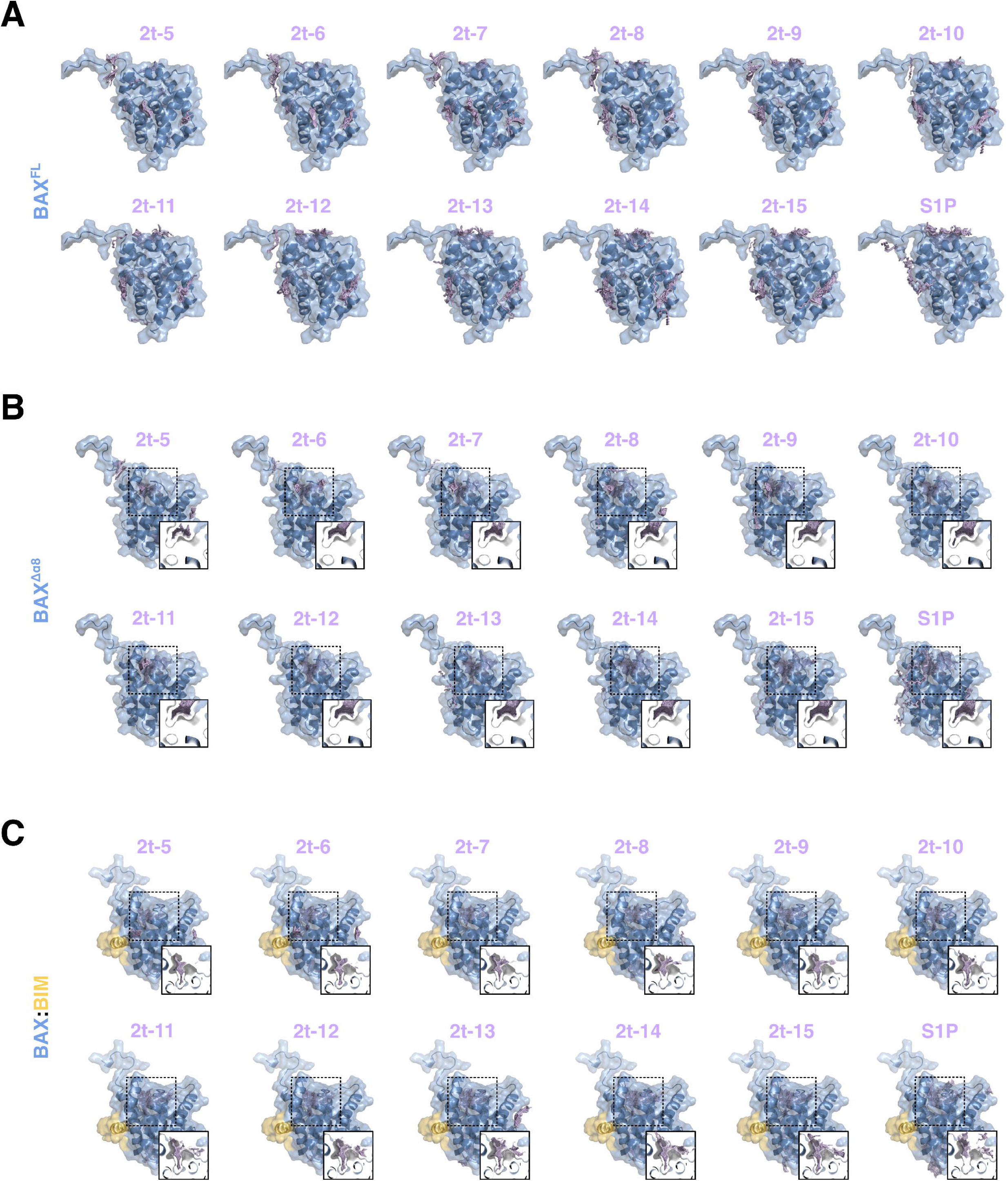
In silico docking simulations suggest long-chain 2t–alkenals exhibit increased specificity for the BAF (Related to Figure 5) **(A–C)** Unbiased *in silico* rigid docking of 2t–alkenals or S1P against BAX using the SwissDock web service. Insets display cross-section of ligands docking within the hydrophobic funnel. Models visualized with PyMOL. **(A)** Visualization of results for each 2t–alkenal or S1P docking on an unmodified structure of BAX (PDB: 1F16, BAX^FL^). Quantification of results summarized in Figure 6C. **(B)** As in **A** with ligands docking on a structure of BAX with alpha helix 8 removed (PDB: 1F16, Δ157–163; BAX^Δα8^). Quantification of results summarized in Figure 6C. **(C)** As in **B** with ligands docking on a structure of BAX bound to a BIM–BH3 peptide with alpha helix 8 removed (PDB: 2K7W, Δ157–163; BAX:BIM).

**Supplementary Figure 7:**
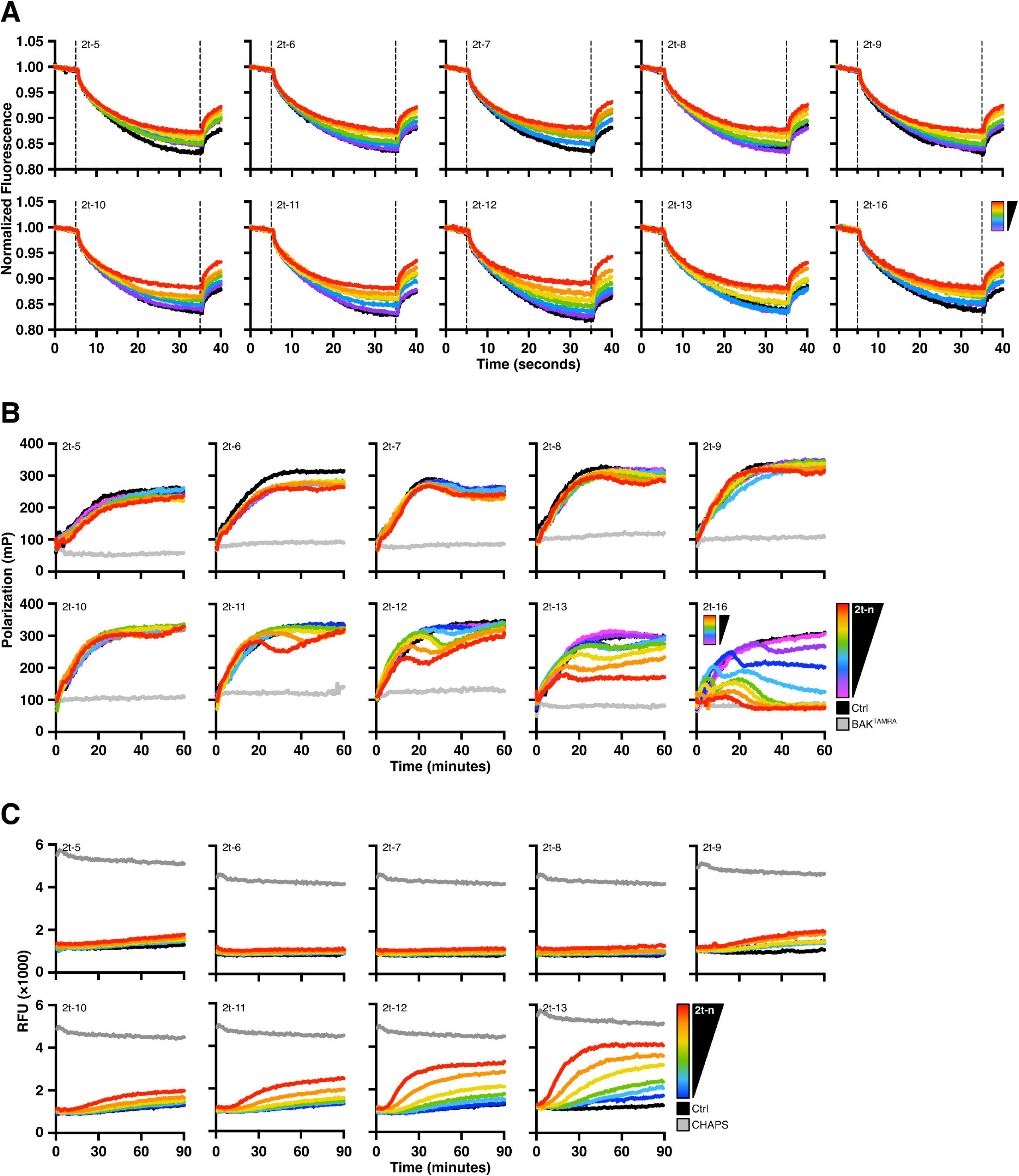
Long-chain 2t–alkenals are capable of BAX activation (Related to Figure 5) **(A)** Alexa Fluor 647-labeled recombinant BAX^WT^ (1 nM) was incubated with CHAPS (0.002%) to inhibit oligomerization, treated with the indicated 2t–alkenals (0.16–5 μM), and subjected to MST. Timetrace data are shown as the mean of replicates. Thermophoresis metrics for each 2t–alkenal are summarized in Figure 5C. **(B)** BAX^2S^ (60 nM) was treated with the indicated 2t–alkenal (3–50 μM), combined with BAK^TAMRA^ (50 nM), and subjected to FLAMBE. Data are shown as the mean of replicates. Parameterized data reporting EP and Tmax for each experiment are provided in Figures 5D**–E**. **(C)** LUVs permeabilized by BAX^2S^ (100 nM) treated with the indicated 2t–alkenal (6.5–50 μM). Data are shown as the mean of replicates. Normalized endpoint permeabilization data summarized in Figure 5F.

**Supplementary Figure 8:**
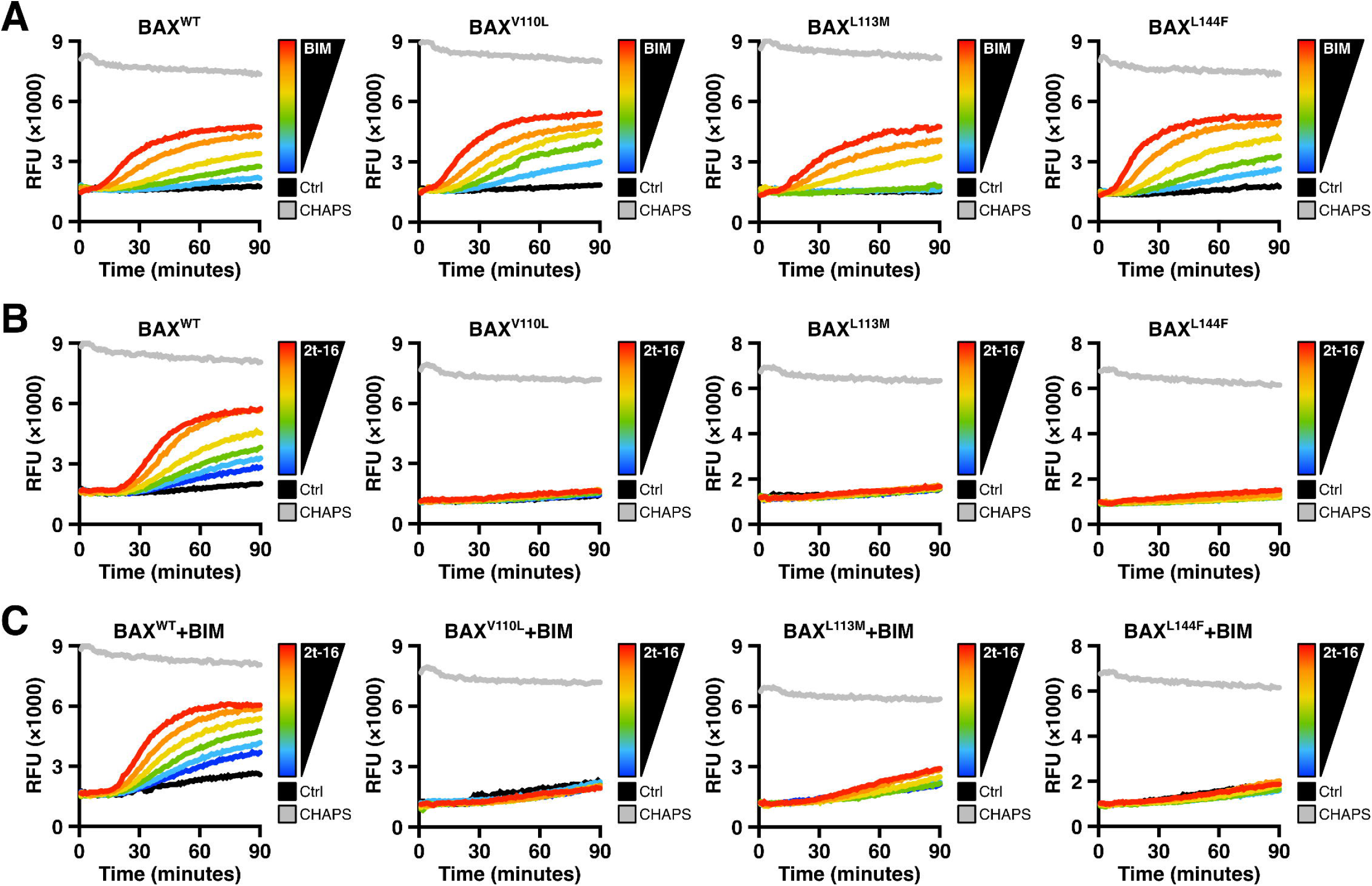
BAF mutations ablate 2t–hexadecenal mediated membrane permeabilization (Related to Figure 6) **(A)** Data summarized by Figure 6G. LUVs permeabilized by WT or BAF mutant BAX activated by BIM–BH3 (0.125−2 μM). **(B)** Data summarized by Figure 6H. LUVs permeabilized by WT or BAF mutant BAX activated by 2t–16 (6.5−50 μM). **(C)** Data summarized by Figure 6H. As in **B** with BAX primed with BIM–BH3 peptide (0.15 μM).

**Supplementary Figure 9:**
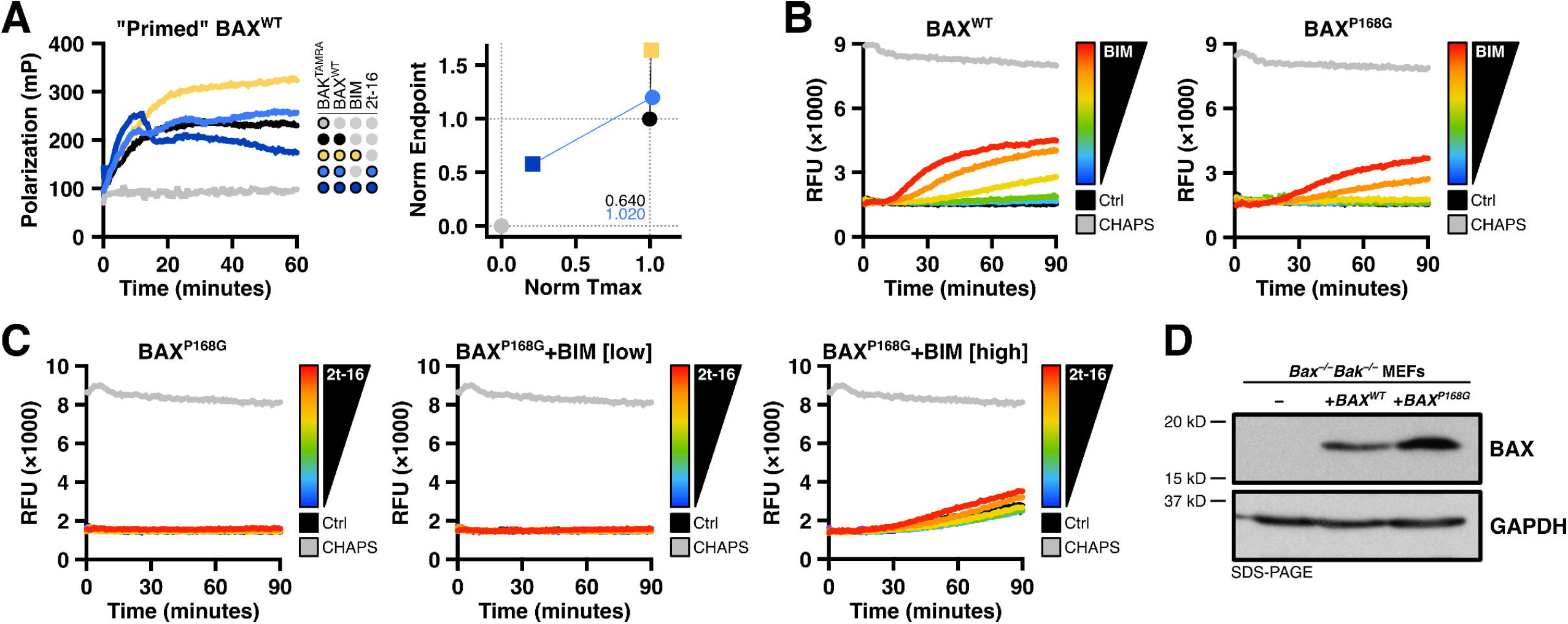
Mutating proline 168 disrupts 2t–hexadecenal synergy with BAX and BIM (Related to Figure 7) **(A)** Left: BAX^WT^ (60 nM) was combined with a non-activating concentration of BIM–BH3 peptide (0.15 μM) and 2t–16 (3 μM), followed by BAK^TAMRA^ (50 nM), and subjected to FLAMBE. Right: Parameterized FLAMBE data in the absence or presence of BIM–BH3 (circle and square datapoints, respectively). Annotations report the magnitude of shift between data with and without BIM–BH3. The parameterized trendline for BAX^WT^ is included for comparison in Figure 7D. **(B)** Data summarized by Figure 7F. LUVs permeabilized by WT or P168G BAX activated by BIM–BH3 (0.125−2 μM). **(C)** Data summarized by Figure 7G. LUVs permeabilized by BAX^P168G^ (100 nM) treated with 2t–16 (6.5–50 μM) ± BIM–BH3 peptide (0.5, 2 μM). **(D)** Western blot confirming expression of BAX protein in transduced *Bax^−/−^Bak^−/−^* MEFs.

## Notes

### Competing Interest Statement

The authors have declared no competing interest.

